# Error, noise and bias in *de novo* transcriptome assemblies

**DOI:** 10.1101/585745

**Authors:** Adam H. Freedman, Michele Clamp, Timothy B. Sackton

## Abstract

*De novo* transcriptome assembly is a powerful tool, widely used over the last decade for making evolutionary inferences. However, it relies on two implicit assumptions: that the assembled transcriptome is an unbiased representation of the underlying expressed transcriptome, and that expression estimates from the assembly are good, if noisy approximations of the relative abundance of expressed transcripts. Using publicly available data for model organisms, we demonstrate that, across assembly algorithms and data sets, these assumptions are consistently violated. Bias exists at the nucleotide level, with genotyping error rates ranging from 30-83%. As a result, diversity is underestimated in transcriptome assemblies, with consistent under-estimation of heterozygosity in all but the most inbred samples. Even at the gene level, expression estimates show wide deviations from map-to-reference estimates, and positive bias at lower expression levels. Standard filtering of transcriptome assemblies improves the robustness of gene expression estimates but leads to the loss of a meaningful number of protein-coding genes, including many that are highly expressed. We demonstrate a computational method, length-rescaled CPM, to partly alleviate noise and bias in expression estimates. Researchers should consider ways to minimize the impact of bias in transcriptome assemblies.

## 1. INTRODUCTION

The development of computational methods for *de novo* transcriptome assembly (Birol et al., 2009; Grabherr et al., 2011a) has empowered a wide array of evolutionary analyses in the absence of a reference genome. Their application has led to significant advances in fields as diverse as phylogenetics (Schwentner, Combosch, Pakes Nelson, & Giribet, 2017), evolutionary responses to climate change (Veilleux et al., 2015), the molecular basis of convergent evolution (Berens, Hunt, & Toth, 2015; Pankey, Minin, Imholte, Suchard, & Oakley, 2014; Velotta et al., 2017), speciation (K. Li et al., 2016), and local adaptation (De Wit & Palumbi, 2013).

All studies using *de novo* transcriptome assemblies rely on two implicit assumptions: that the assembled transcriptome represents an unbiased, if incomplete, representation of the true underlying expressed transcriptome, and that the expression estimates from the assembly are good, if noisy, approximations of the relative abundance of expressed transcripts. Indirect evidence suggests that violations of these assumptions may be frequent. For example, comparisons of different assembly algorithms indicate that there is a degree of complementarity in missingness—one assembler will recover a gene that another cannot, motivating methods to combine assemblies, e.g. the Oyster River Protocol (MacManes, 2018). In other words, particular assemblers may not be able to assemble particular kinds of genes. A separate issue, related to the robustness of expression estimates, is that highly fragmented assemblies comprised of hundreds of thousands of relatively short contigs are pervasive in the literature, presumably representing transcript fragments, haplotypes erroneously designated as separate transcripts, or both. Intuitively, the challenge of mapping reads to fragmented gene models should diminish the robustness of expression estimates. Unknown is whether the compositional aspects of transcriptome assemblies negatively impact SNP calls, allele frequency estimates, and other population genetic parameters.

Below, we evaluate the assumptions inherent to *de novo* transcriptome assembly, discuss how their violation might distort evolutionary inferences, identify particular features of transcriptome assemblies that should be considered during the study design phase of a project, and demonstrate a method for correcting bias in expression estimates. While phylogenomic studies can discard the majority of contigs using aggressive filtering schemes, these are less appropriate for studies that seek to understand patterns of genetic variation or gene expression across populations or closely related species. Therefore, we focus on methodological considerations for this class of investigations.

## 2. MATERIALS AND METHODS

### 2.1 RNA-seq data

Data sets include the *Mus musculus* (C57BL/6J) dendritic cell data used in the original TRINITY paper (MDC), a pool of six whole brains from albino inbred *Mus* (BALB/c), and 8-sample pools of whole brain from wild *M. musculus domesticus* from Massif Central, France (FRA), Iran (IRN), Kazakhstan (KZK), and Germany (DEU). Brain tissue is both transcriptionally complex and expected to express a large proportion of an organism’s overall transcriptional profile. It thus represents a challenging test case for *de novo* transcriptome assembly. Illumina reads were pre-processed using a standard pipeline (Methods S1).

### 2.2 *De novo* transcriptome assembly

We built *de novo* transcriptome assemblies with Trinity v. 2.1.1 (Grabherr et al., 2011b) using default settings. For the *Mus* data sets, to evaluate the robustness of some key findings to the algorithm employed, we also built assemblies using two alternate assemblers: Shannon v. 0.0.2 (Kannan, Hui, Mazooji, Pachter, & Tse, 2016) and BinPacker v. 1.0 (Liu et al., 2016). Where strand-specific reads were available, we ran Trinity and BinPacker with the appropriate strand-specific options. Because the strand-specific mode of Shannon is in experimental phase (Kannan, personal communication), we treated reads as un-stranded in all Shannon assemblies. We assessed assembly quality using standard, commonly used metrics for a) coverage of conserved orthologs to estimate recovery of conserved genes that should be expressed in many tissues, b) the proportion of reads that are aligned to the assembly in a concordant fashion, i.e. whether the assembly is well-supported by the reads and c) assembly contiguity metrics, namely N50,which captures the extent to which the assembly is fragmented (Methods S2). Downstream analyses compare outputs of *de novo* transcriptome assemblies to results obtained from mapping RNA-seq reads directly to the reference genome and its associated annotations of exons, transcripts and genes. (Methods S3).

### 2.3 Assembly and Read Functional Composition

For each contig within a *de novo* assembly, we quantified the proportion of unique assembly nucleotides that were CDS, UTR, intronic, start or stop codon, noncoding elements (including pseudogenic transcripts of genes with “polymorphic_pseudogene” gene biotype), introns of non-coding transcripts, and intergenic/unnanotated sequence. We took a similar approach with sequence reads to determine if the functional composition of RNA-seq reads differed from that of the transcriptome assemblies. For details, see Methods S4.

### 2.4 Expression Estimation

For nine combinations of assembly method (Trinity, Shannon, BinPacker) and *Mus* sample (MDC, BALB/c, Massif Central), we generated expression estimates with RSEM v. 1.2.22 (B. Li & Dewey, 2011a) and Bowtie2, v. 2.2.4 (Langmead & Salzberg, 2012), using RSEM default parameters (which internally sets Bowtie2 arguments). For libraries that were strand-specific, we set the appropriate flags so strand-specificity could be leveraged during the read-mapping stage. We also estimated “true” (reference-based) expression by mapping reads directly onto transcripts extracted from the reference genome, using the same RSEM/Bowtie2 pipeline but with the ENSEMBL GTF annotation.

To be able to evaluate the robustness of expression patterns derived from *de novo* transcriptome assemblies, one must connect assembly contigs to the underlying reference isoforms from which they originated. Thus, after extracting reference transcripts from the genome annotation with RSEM v. 1.2.22 (B. Li & Dewey, 2011b), we mapped transcriptome assembly contigs to these transcripts using BLAT v. 3.1.6 (Kent, 2002). We required at least 95% identity in the alignment to consider a hit valid, but did not set a minimum alignment length in the query (contig). Given the minimal evolutionary distance between query and target sequences, and because we required a minimum of 95% identity for alignments, we did not penalize coverage for mismatches. For each *de novo* assembly contig with > 1 BLAT hit, we identified the true transcript of origin as that for which the alignment produced the maximal coverage. In this way, each contig was uniquely assigned to a single reference isoform.

We assessed concordance between *de novo* assembly and referenced-derived approaches in multiple ways. First, we compared estimates for reference isoforms and their best hit BLAT contig in units of counts per million (CPM), transcripts per million (TPM), and a rescaled CPM that adjusts for differences in effective and observed length estimates between methods (see below, hereafter rescaled CPM). Second, we evaluated concordance between reference gene expression and our best estimate of gene-level expression derived from the transcriptome assembly, by treating all contigs with a best BLAT hit to transcripts of a particular ENSEMBL gene as transcripts of that gene. Contigs without BLAT hits were aggregated by TRINITY components, i.e. “genes”. For Shannon assemblies, the collection of contigs assembled from an individual graph do not necessarily belong to the same gene (Kannan, personal communication). *De novo* assembly gene-level expression was then calculated by building an RSEM index with the *– transcript-to-gene-map* flag, such that contig expression values were aggregated to gene labels specified in a table of gene-contig relationships. Genes identified as expressed in the map-to-reference approach but that did not have any associated *de novo* transcriptome assembly contigs were considered missing.

We also compared gene-level expression analyses between map-to-reference and map-to-*de novo* using expression values estimated with Kallisto v. 0.43.1 (Bray, Pimentel, Melsted, & Pachter, 2016). Because Kallisto operates on the transcript/contig level, we generated gene-level estimates by aggregating to the gene level using the R package Tximport (Soneson, Love, & Robinson, 2015).

We also evaluated how a typical filtering approach influenced the robustness of expression estimates at the gene level. We employed three filtering steps. First, we discarded all lowly expressed transcriptome assembly contigs with TPM ≤ 1. Second, we used Transdecoder v. 3.0.0 (https://github.com/TransDecoder/TransDecoder/wiki) to predict contig peptide sequences, performed BLASTP hit against Uniref90 (downloaded 2017-04-05), and only retained contigs with an alignment e-value ≤ 1 × 10^-5^. Finally, we restricted comparisons between *de novo* transcriptome assemblies and our map-to-reference benchmark to protein coding genes for which at least one isoform was ≥ 200bp; because the default minimum contig size for Trinity is 200bp, the minimum length requirement prevents artefactual downward bias in expression estimates.

The fragmentation of transcript and gene models in assemblies will affect effective length, where the effective length of a transcript equals the length of the transcript minus the mean sequence read fragment length, plus one. In other words, effective length estimates the total number of possible start sites for read alignments to a sequence given the fragment length distribution of reads. For example, if a 1kb transcript is broken into two 500bp contigs in a transcriptome assembly, and the mean read fragment length is 200bp, the effective length of the original transcript equals 801, whereas the summed effective lengths of the two transcriptome assembly contigs equals 602. The observed length also contributes to abundance estimates, e.g. if half of an expressed transcript is not assembled, no counts can be tallied for the missing transcript interval. Because both effective and observed length directly impacts abundance measurements, we assessed whether correcting for differences in both length metrics between reference transcripts and *de novo* assembled contigs would improve concordance in expression estimates. We did this by rescaling CPM as follows:

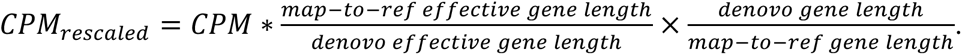

The first scalar adjusts for the difference in estimated effective length, while the second scalar corrects for the difference in observed length. We only evaluated reference features to which at least one transcriptome assembly contig could be aligned.

If biases in expression estimates derived from *de novo* transcriptome assemblies are correlated across samples, this would mitigate downstream effects on differential expression testing. We examined the distribution of differences in such biases among samples, measured as effect sizes. Bias can be quantified as:

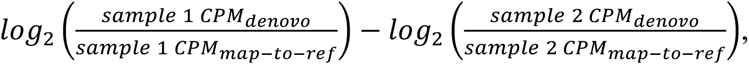

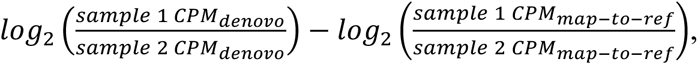

which is the bias observed when comparing log-fold change derived from different methods. We can estimate an effect size as:

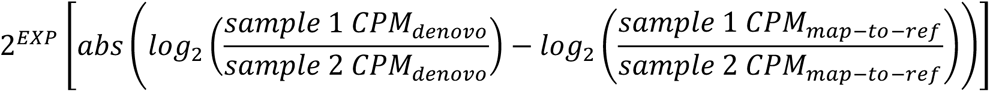

For example, given *de novo* CPM values for samples 1 and 2 of 15 and 20, and for map-to-reference, 5 and 10, respectively, we would estimate that the magnitude of difference between samples would be 1.5 times as great as if one had used the map-to-reference approach.

Two standard ways of filtering a *de novo* transcriptome assembly are to discard contigs that are lowly expressed or that cannot be annotated. For the latter, the usual practice is to find open reading frames, and use BLASTX or BLASTP against a peptide database. We investigated how such filtering impacts the robustness of gene-level expression estimates by recalculating expression estimates after removing contigs with TPM < 1, contigs for which no ORF was detected, and contigs for which a translated ORF did not return a BLASTP hit to Uniref90. We retained the original transcript and gene assignments for assembly contigs derived from BLAT, such that protein hits are simply treated as filter. This approach is optimistic given that it circumvents possible annotation errors, and also due to the fact that *Mus* proteins are prevalent in Uniprot. Because filtering will shorten the total length of sequence that is used to infer expression estimates, this may lead to noisier expression estimates. Thus, we quantified the reduction in coverage of reference annotations induced by our filtering scheme and related it to expression estimates. For details on coverage calculations and filtering, see Methods S5.

### 2.5 Evaluation of variant calls

We compared the map-to-reference genotypes—those derived from mapping reads directly to the genome—to those derived from *de novo* Trinity assemblies. We called variants off of the reference genome using the GATK v. 4 (McKenna et al., 2010), following the Broad Institute’s best practices for RNA-seq data (https://software.broadinstitute.org/gatk/documentation/article.php?id=3891; last updated 2017-01-21). For *de novo* Trinity assemblies, we first constructed SuperTranscripts, (Davidson, Hawkins, & Oshlack, 2017), which attempt to collapse overlapping contigs within Trinity “genes”, and then called variants with the GATK pipeline, based upon python scripts provided with Trinity v. 2.6.5 (see 16ploidy_run_variant_calling_gatk4.py, in the genotying directory of the below-referenced Github repository, which as an example, has the –ploidy argument set to 16 as would be used for the wild *Mus* pools). SuperTranscript alignments to the genome were used to define unique polymorphic site ids that included cases where (a) both methods called genotypes, (b) only map-to-reference called a genotype (including sites with and without valid SuperTranscript alignments), or (c) a genotype was only called off a SuperTranscript. We recorded cases where multiple SuperTranscripts were aligned to the same genomic position, taking the union of all reported alleles as the ST genotype for that position. We restricted our comparison of the two methods by filtering out map-to-reference sites that were indels or contained > 2 alleles. Using genotype calls based upon the reference annotation as the gold standard, we calculated 11 different metrics, defined as follows:

- **Error rate:** # positions with a genotype call in either map-to-reference or map-to-SuperTranscripts, where SuperTranscript alleles do not match map-to-reference alleles ÷ total # positions with a genotype call in either map-to-reference or map-to-SuperTrasncripts; includes positions with no overlapping ST.
- **Error-Snv2Indel**: # biallelic map-to-reference SNVs called as an indel in map-to-SuperTranscripts ÷ # bi-allelic map-to-reference SNVs.
- **FN**: # map-to-reference polymorphic sites with an invariant or missing genotype in map-to-SuperTranscripts ÷ # map-to-reference polymorphic sites; includes sites with no overlapping SuperTranscripts.
- **FN-FractionNoAlignment**: # FN with no aligned SuperTranscript ÷ # FN
- **FN-withAlignment**: Same as **FN**, but restricted to sites with ≥1 overlapping SuperTranscript alignments
- **FP**: # map-to-reference invariant sites with a polymorphic map-to-SuperTranscript genotype ÷ # map-to-reference invariant sites.
- **FP-indel**: # map-to-SuperTranscript indel polymorphic sites at map-to-reference invariant sites ÷ # false positive sites.
- **FP-hetSNV**: # false positive sites with map-to-reference biallelic SNV genotypes ÷ # false positive sites.
- **Recall**: # polymorphic map-to-reference genotypes with matching map-to-SuperTranscripts genotypes ÷ # polymorphic map-to-reference genotypes
- **Recall-withAlignment**: same as **Recall** restricted to sites with ≥1 overlapping SuperTranscript alignments
- **Precision:** # polymorphic map-to-SuperTranscript genotypes that match map-to-reference genotypes ÷ # polymorphic map-to-SuperTranscript genotypes.

In order to determine if SuperTranscripts with either high false negative, false positive or overall genotyping error rates were enriched for particular functions, we performed statistical enrichment tests for Biological Process, Molecular Function and Cellular Component Gene Ontology categories using Panther (Mi, Muruganujan, Casagrande, & Thomas, 2013). For these tests, we assigned SuperTranscripts to gene annotations based upon sequence alignment, and calculated mean SuperTranscript error rate per gene as scores for statistical testing. See Methods S5 for more details.

### 2.6 Code Availability

Software command lines, scripts and accessory files used to implement analyses described above are available at https://github.com/harvardinformatics/TranscriptomeAssemblyEvaluation.

## 3. RESULTS

### 3.1 “High quality” assemblies are highly fragmented

Our assemblies were highly fragmented, consisting of anywhere from 70,375 – 582,133 contigs, with median contig sizes ranging from just a few hundred bases to slightly over 1kb, and N_50_ ranging from 870 to 3021 bp (Table S2). This is typical of many published transcriptome assemblies (Bentley, Haas, Tedeschi, & Berry, 2017; Faherty, Villanueva-Cañas, Blanco, Albà, & Yoder, 2018; Nespolo et al., 2018). Yet, typically employed measures of accuracy and completeness indicate that all assemblies were of reasonably high quality (Results S1, Tables S3 and S4). The large number of short contigs strongly suggest that *de novo* transcriptome assemblies do not typically assembly full length transcripts.

### 3.2 Assembly functional composition is biased

Large fractions of *de novo* transcriptome contigs are dominated by intronic, UTR, and intergenic sequence (Figs S1 and S2). The excess of intronic sequence in contigs is not reflected in the composition of reads (Figs. S1 and S2). The prevalence of intronic and intergenic sequence in assembly contigs (Fig. S3) indicates that a substantial proportion of assembled bases are derived from some combination of transcriptional read-through across exon junctions, assembly of low-abundance pre-mRNA, and low levels of spurious genome-wide transcription.

### 3.3 Frequent genotyping errors

In principle, redundancy through assembly of haplotypes as distinct contigs should lead to erroneous genotype calls and biased estimates of allele frequency spectra. For example, two contigs can actually be haplotypes of a single locus, such that a truly heterozygous position is instead represented as two separate contigs, one homozygous for the reference allele and one homozygous for the alternative allele. Two lines of evidence indicate such redundancy is present in transcriptome assemblies. First, BLAT all-by-all alignments indicate that many contigs are nearly identical (Figs. S4-S6), in some cases differing only by single nucleotide variants (e.g. Figure S4 B). Surprisingly, Trinity assemblies will produce such highly similar contigs from different assembly graphs which are purported to assemble isoforms of a gene or sets of closely related paralogs (Fig. S6). Second, there is a high frequency of cases where all nucleotides of a transcript from a single-isoform *Mus* gene are covered by ≥ 1 *de novo* assembly contig (Fig. S7). The latter are primarily due to slight variations in the assembly of intronic bases, and secondarily due to differences in CDS content (Results S2, Fig. S8).

The opposite problem can happen when paralogous loci are collapsed, leading to a heterozygous genotype call where there would otherwise be two invariant sites, or perhaps a mixture of alleles due to, for example, collapsing of an invariant and a heterozygous position. To assess the degree to which these and other errors affect transcriptome assemblies, we compared genotype calls and derived diversity metrics between the map-to-reference and map-to-transcriptome approaches using 11 different metrics.

The frequency of errors, in which the allele set recovered by the two methods differed or in which a genomic position with a called genotype contained no overlapping SuperTranscript nucleotides, was high, ranging from 30-37% for the wild samples, and 49-83% for the inbred samples (Table 1). Error rates varied among SuperTranscripts within a *de novo* transcriptome assembly, with some SuperTranscripts demonstrating extremely high error rates. While the median error rate per SuperTranscript was 0.14% per base or less, maximum per SuperTranscript rates typically exceeded 10% per base (Table S5). For five of six samples, the median number of errors per SuperTranscript was 2, but, depending upon the sample, as many as 56-187 errors could occur within a single SuperTranscript (Table S5). Thus, within any *de novo* transcriptome assembly, a small number of contigs may contribute disproportionately to noise that may obscure the true patterns of polymorphism. On a per-sample basis, errors show a modest yet statistically significant bias towards the distal end of a SuperTranscript for all samples except MDC (sample-level Wilcoxon 1-sided tests, H_0_ =0.5, p < 2 × 10^-16^; Fig. S9).

**Table 1.** Selected performance metrics comparing genotypes derived from SuperTranscript collapse of TRINITY de novo transcriptome assemblies with those derived from a map-to-reference genome approach. Samples are *Mus dendritic cell* (MDC), albino inbred mouse (Balb/c), and wild mouse from France (FRA), Germany (DEU), Iran (IRN), and Kazakhstan (KZK). See **MATERIALS AND METHODS** for information on metric calculations.

| Metric | Sample |  |  |  |  |  |
| --- | --- | --- | --- | --- | --- | --- |
|  | MDC | BALB/c | FRA | DEU | IRN | KZK |
| Error rate | 0.829 | 0.494 | 0.350 | 0.367 | 0.327 | 0.296 |
| Error-Snv2Indel | 0.194 | 0.160 | 0.282 | 0.307 | 0.281 | 0.248 |
| False negative (FN) | 0.558 | 0.712 | 0.244 | 0.245 | 0.228 | 0.255 |
| FN-withAlignment | 0.371 | 0.587 | 0.145 | 0.144 | 0.134 | 0.150 |
| FN-FractionNoAlignment | 0.532 | 0.424 | 0.474 | 0.482 | 0.475 | 0.484 |
| False positive (FP) | 0.871 | 0.267 | 0.388 | 0.462 | 0.396 | 0.151 |
| FP-indel | 0.187 | 0.060 | 0.178 | 0.219 | 0.188 | 0.067 |
| FP-hetSNV | 0.621 | 0.191 | 0.195 | 0.225 | 0.193 | 0.077 |
| Recall | 0.442 | 0.287 | 0.754 | 0.752 | 0.769 | 0.742 |
| Recall-withAlignment | 0.628 | 0.410 | 0.852 | 0.853 | 0.862 | 0.846 |
| Precision | 0.145 | 0.169 | 0.780 | 0.762 | 0.814 | 0.774 |

The false negative rate (FN; 23-71%) was greater than the false positive rate (FP; 15-87%) in four of six samples. The latter includes all classes of errors including indel polymorphism. Anywhere from 42-53% of FN were due to there being no SuperTranscript aligned at a genomic position where a map-to-reference heterozygote genotype call was made (Table 1 FN-FractionNoAlignment). Given that rates of mapping SuperTranscripts to the genome were extremely high (>99%), these FN instances are primarily due to a failure to assemble transcripts overlapping polymorphic genomic positions. Even at sites called as heterozygotes in map-to-reference analyses that were assembled into SuperTranscripts, false negative rates were still relatively high, from 13-15% and 37-59% in the wild and inbred samples, respectively (Table 1). False negative rates demonstrated significant negative correlations with SuperTranscript length in five of six samples, but those correlations were weak (Spearman’s ρ, -0.24- -0.10), indicating a minimal increase in false negatives in shorter SuperTranscripts.

Because read depth influences the ability of tools such as GATK to call genotypes, we suspected false negatives might be due to reduced read depth at homologous genomic positions called as heterozygotes with the map-to-reference approach. While map-to-reference and map-to-SuperTranscript coverage were largely colinear for concordant and false positive map-to-SuperTranscript genotypes (Figs. S10 and S11), false negative sites with overlapping SuperTranscripts displayed small reductions in SuperTranscript read depth relative to map-to-reference at sites where SuperTranscripts mapped to the genome (Fig. S12). Given the magnitude of coverage reduction is small, we suspect it may not be the only factor contributing to false negatives at sites where a SuperTranscript covers a polymorphic genomic position.

The false positive (FP) rate ranged from 15-87%. Incorrect heterozygous and indel genotype calls at truly invariant sites are expected when paralogs are collapsed, and both occur with considerable frequency in our samples. From 6-22% of FPs are erroneous indel genotypes, 8-62% of FPs are erroneous heterozygous genotype calls, and the remainder are other classes of SuperTranscript polymorphisms, including both indel and single nucleotide variants, with > 2 alleles. The inference of frequent paralog collapses is also supported by the high rate at which true SNV polymorphisms are called as indels in SuperTranscripts (Error-Snv2Indel, 16-31%). False positive rate was significantly associated with SuperTranscript length in all samples, but showed contrasting signals between inbred (MDC, Balb/c) and wild (FRA, DEU, IRN, KZK) samples: correlations for inbred samples were moderate and negative (Spearman’s ρ, −0.27 to −0.49), but weak and positive for wild samples (Spearman’s ρ, 0.01 to 0.05).

When genotypes were called in SuperTranscripts, the frequency of calls that were correct (i.e. precision) ranged from 66-83% for the wild samples, but precision for the inbred samples, particularly MDC, was substantially lower. Recall ranged from 76-81% in the wild samples; as with precision, recall for both inbred samples was substantially lower.

We examined false negative, false positive, and overall error rates more closely, evaluating whether particular GO terms were enriched in SuperTranscripts with higher error rates. Panther statistical enrichment tests indicated that there were no categories enriched for false negatives, and only a few for false positives in the FRA sample, indicating there is no broader pattern of enrichment for these types of errors (Table S6). With respect to overall error rates, FRA SuperTranscripts with high error rates were enriched for MHC class 1 protein complex genes, which contains hundreds of paralogs, but these terms were not enriched in any other sample (Table S6). Strong signals for gene families were not detected, with the only terms that consistently showed up as being enriched related to ribosomal components (Table S6).

Overall, the frequency of genotyping errors seems likely to distort evolutionary inferences and reduce power across a wide range of statistical methods, particularly those aimed at detecting signals of selection on particular polymorphisms, allele-phenotype and allele frequency-environment associations.

### 3.4 Negatively biased heterozygosity estimates

In all but the inbred strain samples (MDC and BALB/c), we observed reduced heterozygosity in the exonic regions of SuperTranscripts relative to mapping to those regions in the reference genome (Fig. 1). A substantial contributor to this bias is the fact that, as noted above, approximately half of map-to-reference heterozygous positions are not overlapped by any SuperTranscript. In addition, this bias does not result from SuperTranscripts being preferentially assembled in regions of lower heterozygosity, given that, for the wild samples, map-to-reference heterozygosity is actually higher in SuperTranscript-overlapping regions (Fig. S13). Furthermore, the bias cannot be due to some unique feature of genotype calls in exons, as the same pattern holds when considering all genotype calls (Fig. S14). Another possible explanation may be lower power to detect low frequency alternative alleles in map-to-SuperTranscript analyses. However, in our five samples consisting of pooled individuals, minor allele frequency at map-to-reference heterozygous sites did not vary between correct SuperTranscript heterozygous calls, false negatives, and other genotyping errors (Fig. S15). Downward bias in heterozygosity estimates occurs despite a relatively high prevalence of false positive heterozygotes. Because this bias shifts all estimates towards zero, it truncates the observed range of variation, making it more difficult to make meaningful inferences from comparisons of diversity across genes, populations, and species.

**Figure 1.**
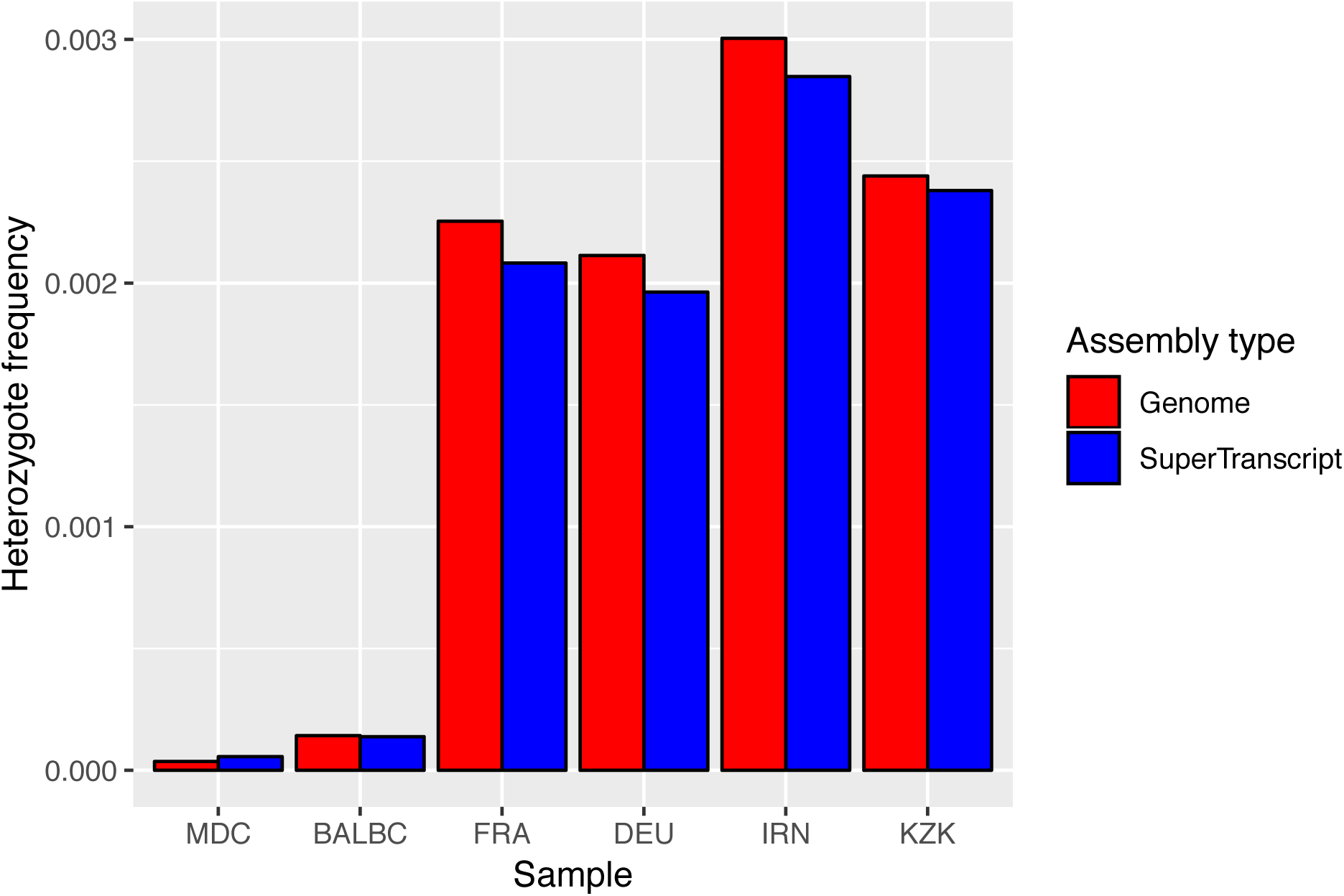
Heterozygosity per base for exonic biallelic SNV variants called relative to the *Mus* reference genome using the GATK best practices pipeline versus those obtained from supertranscripts constructed from TRINITY assemblies using scripts provided with the Trinity v. 2.6.5 package. See **Materials and Methods** for details concerning how exonic coordinates were obtained for SuperTranscripts.

### 3.5 Single contigs are poor expression estimators

Expression estimates derived from the contigs that are the best BLAT hit relative to the reference transcripts are only weakly correlated (Spearman’s ρ: CPM: 0.070-0.387; TPM: 0.045-0.392; rescaled CPM: 0.083-0.415), with expression for those transcripts, and show positive bias for lowly expressed reference transcripts (Fig. S16 A and B, Table S7). Correlations between best-hit contig expression and expression of the associated reference gene, after accounting for effective length differences, are slightly better but still relatively weak (Spearman’s ρ: rescaled CPM: 0.103-0.657, Table S6). Overall, the use of single contigs as proxies for gene-level expression appears problematic.

### 3.6 Correctable gene-level expression bias

Expression estimates derived from *de novo* transcriptome assemblies are far more robust at the gene level, when one is able to group contigs into genes via annotation (Fig. 2, Table S8). It is reassuring that the correlation between the methods is quite strong, although there is sizeable positive bias at lower expression levels for TPM, which is substantially reduced for CPM. Furthermore, for many genes there are substantial deviations from estimates obtained with mapping to the reference, even at high expression levels (Fig. 2, Fig. S7). For estimates generated with RSEM, bias and deviations from the map-to-reference estimates are reduced when one rescales CPMs to account for differences in effective length estimates (Fig. 2C). This indicates that effective length differences—that is, the difference in the number of start sites for read alignments, given a particular sequenced fragment size, between a full-length transcript, and a set of transcript sub-sequences represented by contigs—are part of the problem, and in particular, that effective length is not properly estimated on the fragmented gene models that are typical in *de novo* transcriptome assemblies. Biases in gene expression are reduced as the difference in effective length between map-to-reference and map-to-transcriptome-assembly decreases (Fig. 3). Even so, disparities in effective length estimates between map-to-reference and map-to-*de novo*-transcriptome-assembly approaches can persist even at high reference gene expression, with many *de novo* transcriptome-based estimates of gene expression showing a fourfold or greater deviation relative to the map-to-reference benchmark (Fig. 2, Fig. S18). Correlations between reference gene expression and gene-level summing across assembly contigs obtained with Kallisto were very similar, if slightly inferior, to those obtained with RSEM (Results S3, Table S8, Fig. S17).

**Figure 2.**
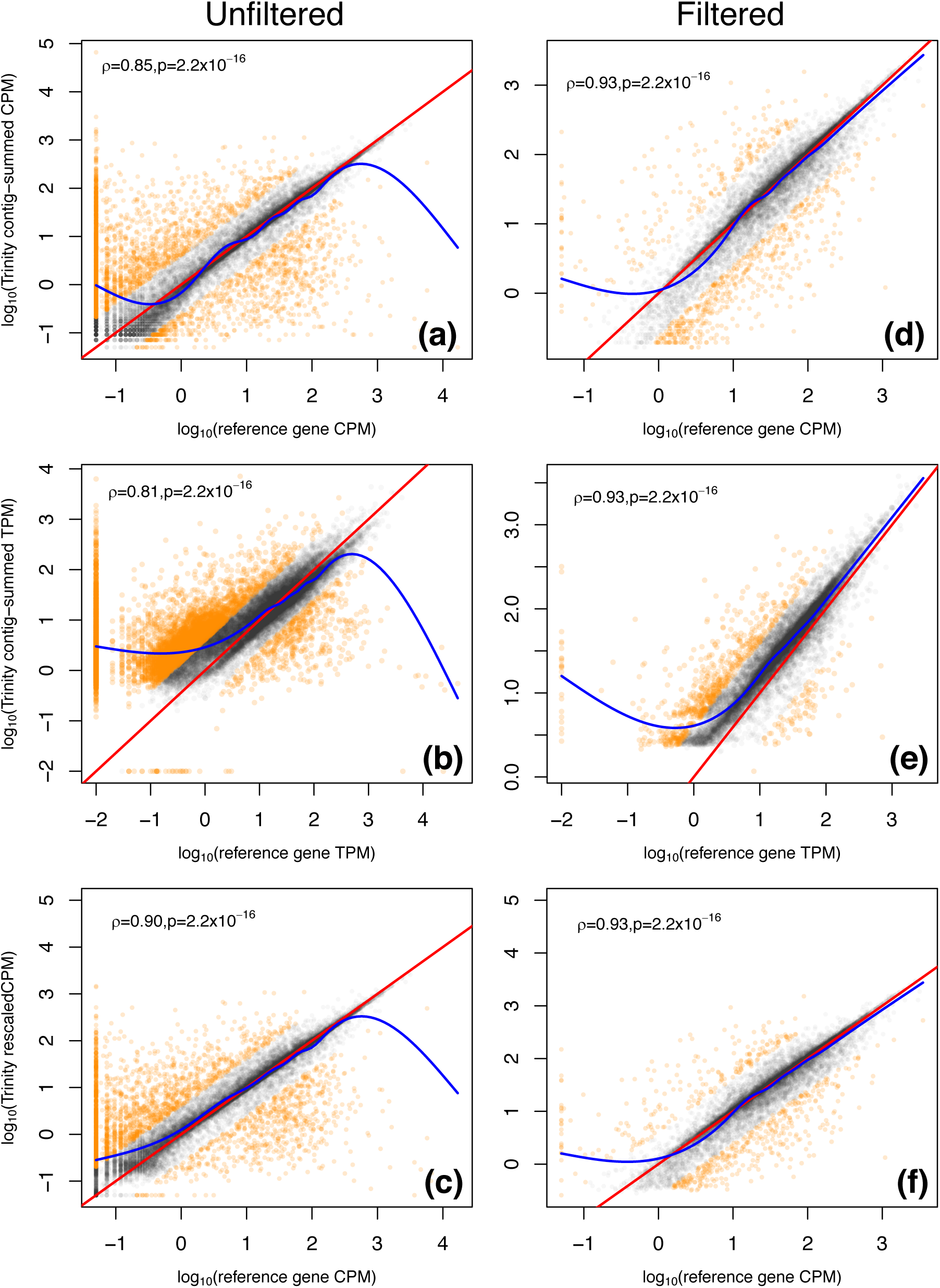
For the FRA data set, comparison of gene-level expression summed across BLAT alignments of TRINITY contigs to reference transcripts (grouped by gene), to reference gene expression in terms of **(a, b)** CPM, **(b, e)** TPM, and **(c, f)** rescaled CPM that accounts for differences in effective length estimates between methods. Left and right columns are for unfiltered and filtered transcriptome assemblies, respectively. For CPM, TPM, and rescaled CPM, 0.05, 0.01 and 0.05 are added to values prior to log-transformation, in order to visualize instances where expression = 0 for one of the methods being compared. The red line represents unity, the blue line is the smoothed fit to data. Gray and orange points indicate log-fold change of Trinity/map-to-reference genes less than and ≥ 2, respectively. We exclude all cases where an expressed reference gene has no TRINITY contigs aligned to it, such that it is missing from the *de novo* transcriptome assembly.

**Figure 3.**
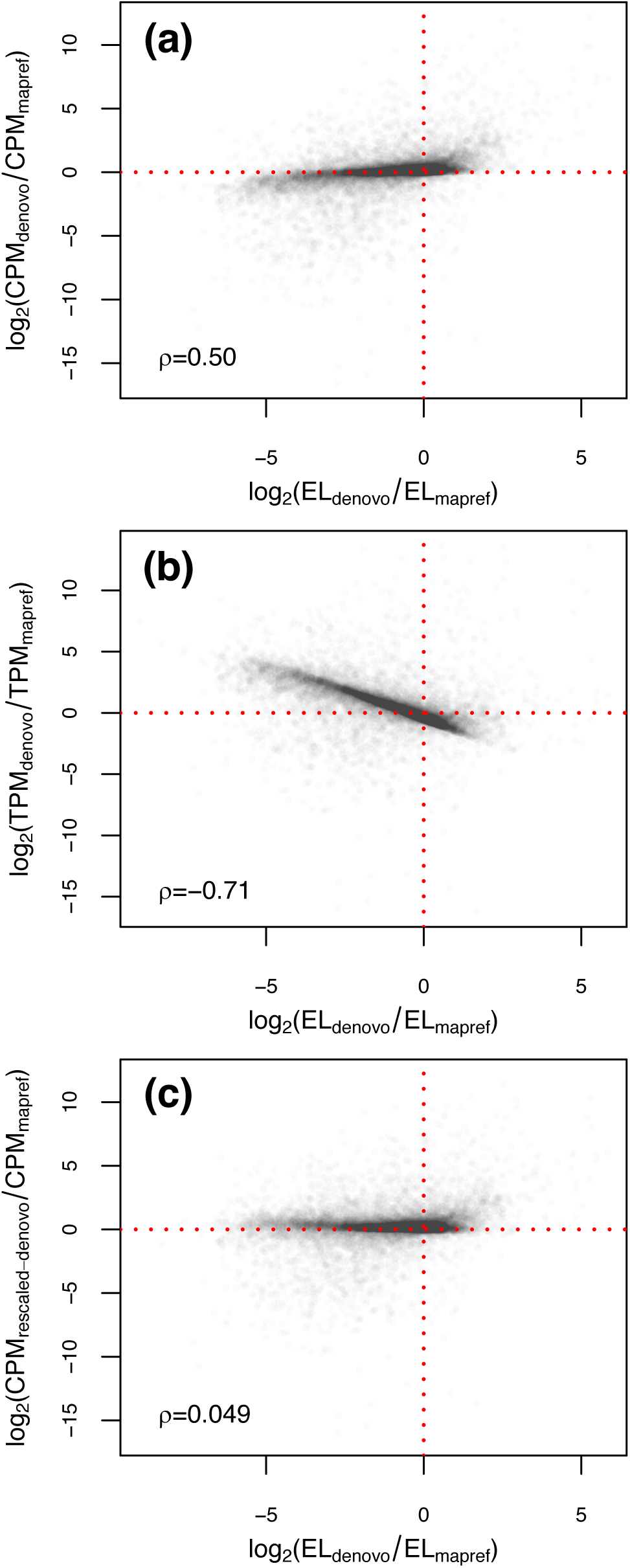
For the FRA data set, for genes with non-zero expression in both transcriptome assembly and map-to-reference, log-fold change between TRINITY and map-to-reference expression estimates as a function of difference in effective length (EL) for **(a)** CPM, **(b)** TPM, and **(c)** rescaled CPM.

### 3.7 Expression bias and DE testing

Biases in expression estimates derived from transcriptome assemblies might not have large effects on downstream differential expression analyses if biases are consistent across samples. While the pooled *Mus* data sets we used did not permit us to look at inter-individual variation amongst individual libraries used to build a single assembly, we looked across the wild *Mus* datasets to see if biases in gene-level expression were concordant, i.e. were biases for a particular gene always of similar magnitude and in the same direction. We focus on results for Trinity assemblies of wild *Mus* samples, as Trinity consistently showed stronger correlations with map-to-reference based estimates, such that our findings are conservative with respect to assemblers showing weaker correlations. Reassuringly, log-fold change of transcriptome assembly vs. map-to-reference expression estimators (LFC_METHOD_) showed broad patterns of moderate concordance at the gene level indicating that gene-level features influence the magnitude of bias (Fig. S19). Inter-sample correlations were, for CPM, 0.625-0.727; TPM, 0.790-0.830; and rescaled CPM, 0.605-0.671. These correlations are weaker for CPM and rescaled CPM, in part, because biases are reduced overall, compressing LFC_METHOD_ toward zero. Despite these broad correlations, a number of genes may show excessive bias. We measured this as a bias effect size that indicates how much greater a difference in expression would be detected using a *de novo* transcriptome approach, relative to mapping to reference annotation. Comparing the FRA and DEU samples, for CPM, TPM, and rescaled CPM, 5.24%, 5.16% and 3.35% of genes showed bias effect sizes ≥ 5, with more than 2% of genes showing an order of magnitude of difference between methods of ≥ 10 for CPM and TPM (Fig. 4).

**Figure 4.**
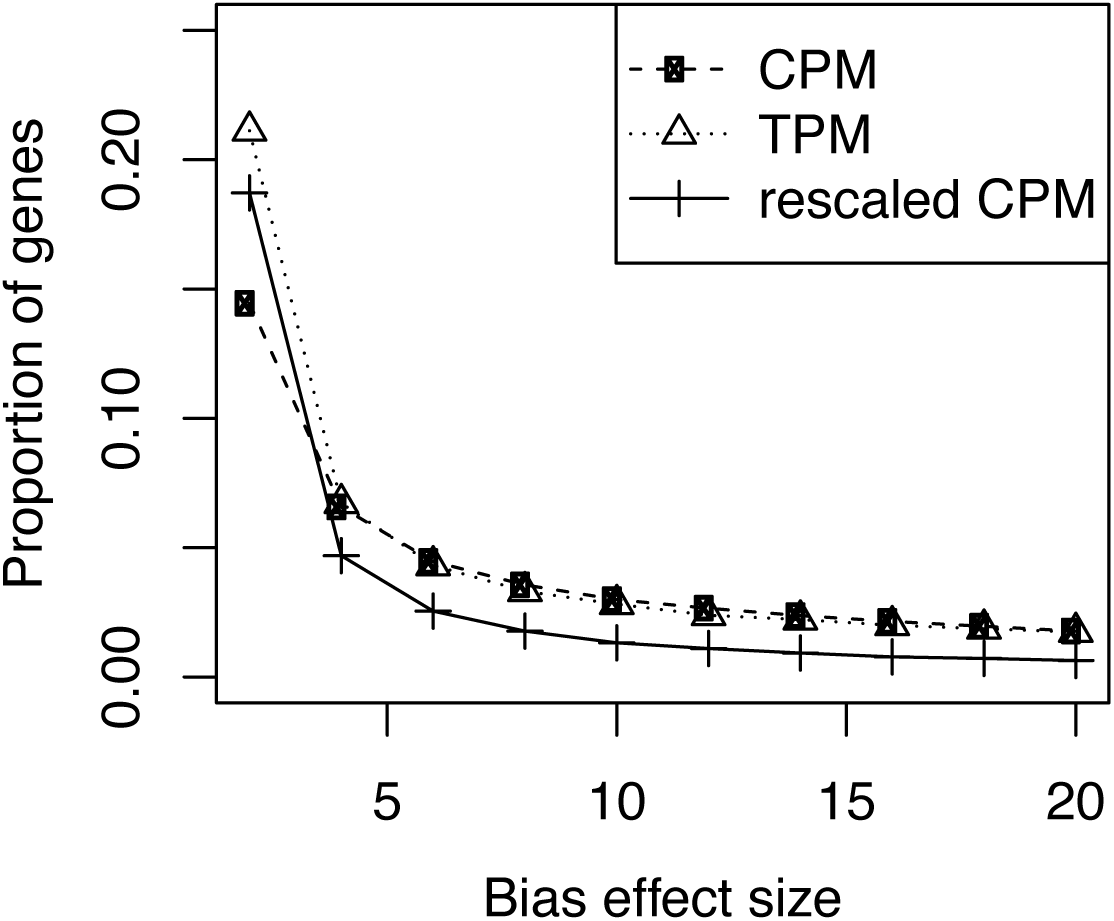
Frequency of bias effect sizes for gene-level expression estimated by CPM, TPM, and rescaled CPM for a TRINITY-based comparison of the FRA and DEU samples.

### 3.8 Effects of filtering on expression estimates

Filtering out lowly expressed *de novo* transcriptome assembly contigs as well as those contigs without a BLASTP hit to Uniref90 led to a noticeable improvement in the correlation between map-to-transriptome assembly and map-to-reference abundance estimates for protein coding genes (Fig. 2 D, E, and F). Post-filtering, bias and noise in TPM was dramatically reduced. However, despite a correlation nearly identical to that for CPM and rescaled CPM, there is consistent positive bias across the range of map-to-reference expression (Fig. 2E). Nevertheless, the improvement across all three metrics relative to unfiltered assemblies belies a tradeoff that is hidden from view from researchers without access to an annotated genome: filtering leads to a loss of many expressed protein coding genes that would be recovered without filtering. For the FRA assembly, 21.0% of such genes are filtered out, although only 10.6% of genes are lost if one excludes from consideration lowly expressed genes with map-to-reference TPM < 1 (Fig. 5). Depending upon the sample-assembler combination, 9.1-32.5% of expressed protein coding genes can be lost due to filtering, and for genes with map-to-reference TPM ≥ 1, 6.0-27.1% (Table S7). In nearly all sample-assembler combinations, the proportion of genes lost due to filtering far exceeded the proportion that were missing due to a failure to assemble those genes, with most exceptions occurring for BinPacker assemblies, which performed poorly with respect to recovery of expressed genes (Table S7).

**Figure 5.**
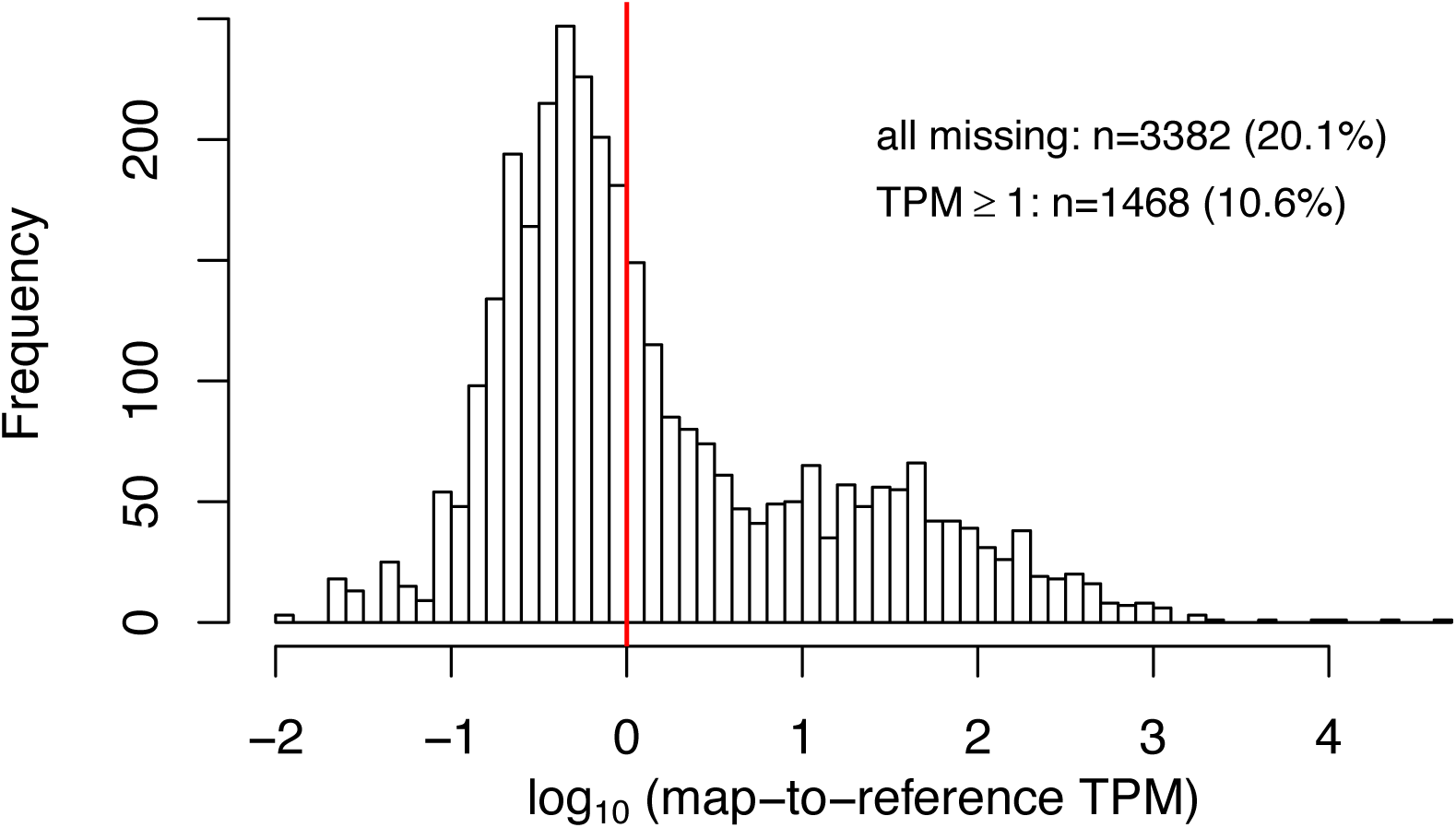
Histogram of map-to-reference TPM for expressed protein-coding genes that were detected in the FRA TRINITY assembly prior to filtering out un-annotated and lowly expressed assembly contigs, but were lost due to filtering.

## 4. DISCUSSION

*De novo* transcriptome assemblies have been an important entry point for functional and population genomics in non-model organisms, but the two key assumptions underlying their application—that the composition of *de novo* transcriptomes is unbiased if incomplete, and that expression estimates derived from them are unbiased if noisy—are consistently violated, albeit in unexpected ways. Below, we explain how particular compositional aspects of *de novo* transcriptome assemblies may impact estimates of gene and isoform expression, genotype calls, and genetic diversity—estimates which are frequently used for making evolutionary inferences for non-model organisms.

SNP calls and allele frequencies inferred from them and that are derived from transcriptome assemblies have been used in a variety of contexts, including estimation genetic diversity in evolutionary analyses (Romiguier et al., 2014), population structure (Tepolt & Palumbi, 2015), reconstruction of demographic history and gene flow (De Wit & Palumbi, 2013; Le Gac et al., 2016; Roda, Mendes, Hahn, & Hopkins, 2017), estimation of dN/dS (Cokus, Gugger, & Sork, 2015), F_ST_ outlier analysis (Bankers et al., 2017), and tests for allele-specific expression (Singh et al., 2018). Given this wide application of *de novo* transcriptome assembly genotypes in studies of non-model organisms, by far the most unexpected findings from our study were the extremely high genotyping error rates derived from transcriptome assemblies, specifically SuperTranscript representations of TRNITY assemblies. False negatives were due, in part, to the absence of a SuperTranscript aligned to a putatively true polymorphic site recovered with a map-to-reference approach. False negatives are due to either the failure to assemble any contigs at all for a gene (i.e. missing genes), or the incomplete coverage of genes not extending over some fraction of polymorphic sites. That genotyping errors were biased towards the distal end of contigs suggest that assembly graphs are getting broken at hard to assemble features containing low complexity and/or increased polymorphism. When such graph-splitting occurs, the intervening polymorphic sites will, by definition, not be covered by a SuperTranscript, leading to downwardly biased estimates of heterozygosity in *de novo* assemblies. False negatives may have also been caused by the slightly reduced read depth at homologous positions in read alignments to SuperTranscripts. That the reduction in read depth, was, on average, small suggests that other factors may play a role. It may be that genotype inference methods such at the GATK Haplotype Caller that leverage haplotypic information, have poorer performance when iterating over alignments to relatively short assembly contigs. Another possibility is that assembly errors may generate patterns that get flagged by standard genotyping filters. In contrast, false positives appear to be driven by paralog collapses given the bias towards heterozygous false positive sites.

While we document overall high error rates, the range of precision we observed was broadly consistent with what was observed in the original paper that describes the LACE algorithm (∼ 80%) for converting transcriptome assemblies into SuperTranscripts (Davidson et al., 2017). It is important to note that, in that paper, recall and false negative rates are not reported, and genotyping performance is restricted to SuperTranscript genotype calls relative to known variants in Genome in a Bottle Consortium high confidence regions for humans (Zook et al., 2016). Such well-curated, filtered benchmark SNV sets are, by definition, not available for non-model organisms with neither genomes nor extensive resequencing data. Results reported on them are optimistic relative to what can be expected for organisms without such genomic resources. For this reason and to create more realistic expectations for molecular ecologists, we use genotypes obtained from mapping to reference genome as our truth set.

The negative bias we observed in heterozygosity estimates derived from *de novo* transcriptome assemblies indicates that the underlying site-frequency spectrum derived from an SuperTranscript assembly will be biased in ways that distort other population genetic summary statistics typically reported in molecular ecological investigations that utilize genotypes called from transcriptome assemblies. More broadly, the error rates we document suggest that several lines of transcriptome-assembly based research should be approached with caution. First, erroneous heterozygous calls, when added to the redundancy problems described above, will substantially reduce power and elevate false positive rates in allele-specific expression analyses. Second, levels of genome-wide patterns of genetic diversity derived from transcriptome assemblies will likely be negatively biased, at least for diploid species. The distinction concerning ploidy is an important one because, in the absence of a genome or karyotypic information, there can be no a priori expectation of how events such as whole-genome or even chromosome-level duplication will impact genotyping inference. Thus, both integration of diversity metrics across transcriptome assemblies with differing, unknown levels of ploidy, and comparing transcriptome and genome assembly-derived metrics will confound method with biology and can lead to spurious inferences. Even when applied solely to diploid transcriptome assemblies, as haplotypes will get assembled into different SuperTranscripts, the expectation is that negative bias will be greater for more heterozygous genomes, such that estimated levels of diversity across assemblies will artificially appear more similar than they are in actuality, impeding researchers’ ability to detect underlying patterns of variation. Overall, the distortion of allele frequencies derived from transcriptome assemblies strongly suggests that they will inject bias into analyses of population structure, demographic inference, and efforts to detect positive selection. Whether or not improved clustering of contigs can mitigate this distortion remains an open question.

Our results clearly indicate that isoform-level analyses and using individual contigs as proxies for isoform or gene-level expression, should be avoided. Even when annotation-based groupings of contigs are performed in an optimal setting, such as our analyses of *Mus* transcriptomes, there is substantial noise in gene-level expression estimates derived from transcriptome assemblies relative to a map-to-reference based approach, and there is positive bias at lower expression levels. Incorrect estimation of effective length metrics used in RNA abundance calculations play a substantial role in the observed patterns of bias. Nevertheless, it may be surprising to researchers that biases persist for standard length-adjusted metrics such as TPM. CPM has less bias compared to TPM. Bias and noise can be reduced by rescaling CPM to account for differences in effective and observed length, such that the rescaled CPM more faithfully recapitulates map-to-reference based expression estimates, even if substantial noise persists. Adjusting expression estimates to account for effective and observed length differences can be beneficially used in two different applications. First, one could conduct differential expression estimates across different transcriptome assemblies when it is necessary to generate different assemblies for different conditions, such as divergent species or sub-species. Because one of the first steps differential expression tools execute is to normalize by library size, one can simply apply the correction directly on the counts themselves. CPM and count scale are directly proportional, with the former effectively being the latter multiplied by a sample-specific constant, such that all benefits we report on rescaled CPM will apply to similarly transformed counts. Rescaling counts across a set of species would require treating one species as the “reference”, and rescaling counts for other species relative to it. Outside of the realm of formal tests of differential expression, we suggest that rescaled CPM could be useful in phylogenetic comparative analyses of expression at the gene level. One of the big challenges of these methods is accounting for length differences between transcript/contig sets belonging to orthologous genes, and the adjusted CPM appears to substantially outperform other metrics in this regard. While our method for correcting for effective length differences substantially reduced the frequency of bias outliers, it does not eliminate them entirely. Further investigation of other sources of bias is necessary in order to devise methods to mitigate their impacts on expression analyses. Even so, we should note that our adjustment method is relatively simple, and that it may be possible to adopt more sophisticated approaches that leverage such information as sequence composition and complexity for additional improvements.

Standard filtering of *de novo* transcriptome assemblies, including removing lowly expressed and un-annotated contigs, can measurably improve the concordance between transcriptome assembly and map-to-reference-based expression estimates. Unfortunately, filtering leads to the loss of many expressed protein coding genes that are detected in a *de novo* transcriptome assembly prior to filtering. Researchers relying on *de novo* transcriptome assemblies for expression analyses thus face a difficult tradeoff—get less robust expression estimates for more genes, and risk a larger fraction of erroneous differential expression results, or get more robust estimates for far fewer genes, and risk missing those that could be functionally relevant to the phenotypes being investigated.

We do not test here datasets amenable to differential expression analyses, and a counter-argument to our findings of bias and noise is that neither should have a substantial effect, so long as bias and deviations from map-to-reference expectations are consistent across replicates and conditions. While we did find broad congruence in bias across samples, gene-level differences in bias among samples were frequent enough such that they will likely elevate false positive rates for differential expression tests derived from *de novo* transcriptome assemblies. This effect can be minimized by utilizing metrics that adjust for effective length differences.

## 5. CONCLUSIONS

Given the challenges associated with eco-evolutionary analyses that rely on *de novo* transcriptome assemblies and the rapidly falling costs of long read sequencing, an unavoidable recommendation for research teams launching new studies may be to generate a provisional genome assembly, e.g. using a combination of Oxford Nanopore or PacBio long reads and Illumina paired-end short reads (Austin et al., 2017; Marcionetti, Rossier, Bertrand, Litsios, & Salamin, 2018; Michael et al., 2018; Nowoshilow et al., 2018), at least in some cases. Such an assembly can be annotated with *ab initio* gene prediction tools and Illumina RNA-seq paired-end reads. Given a reasonable quality annotated genome, fewer reads per sample would have to be allocated for downstream expression analyses, and the cost savings from lighter RNA sequencing would partially offset costs of genomic sequencing and *de novo* assembly. Nevertheless, there are three reasons whole *de novo* transcriptome assembly might still be attractive. First, genome assembly remains challenging and expensive species with large genomes or with unusual genome structure precluding standard methods. Second, thousands of RNA-seq data sets now exist for species without reference genomes, and it is highly unlikely that genome assemblies will be carried out any time soon for anything but a small fraction of these species. Third, analyses based upon genome assemblies will depend strongly on the quality of the assembly. Even for genomes of reasonably high quality, protein coding genes may be missing, either through problems with the annotation process or due to the fact that genes fall into assembly gaps (Peona, Weissensteiner, & Suh, 2018). In other words, genome assembly is not necessarily a panacea for all problems related to expression analyses. However analytically challenging, there is great potential for employing *de novo* transcriptome assemblies in interesting comparative analyses, so long as the conducted in an informed fashion—avoiding analyses modes that suffer from fundamental biases, mitigating biases where possible for others, and, always, placing appropriate caveats given the biases that will inevitably remain.

## Supporting information

Supplemental methods, results, tables ane figures

## ACKNOWLEDGMENTS

The computations in this paper were run on the Odyssey cluster supported by the FAS Division of Science, Research Computing Group at Harvard University. We thank members of the FAS Informatics Group for discussion. We also thank Nadia Davidson, Alicia Oshlack, Robin Hopkins, and Mara Laslo for feedback on an earlier version of this manuscript. This work was conducted on the traditional territory of the Wampanoag and Massachusett peoples.

## DATA AVAILABILITY

Upon acceptance for publication, all transcriptome assemblies, SuperTranscript assemblies, and vcf genotype files will be made available on Dryad.

## AUTHOR CONTRIBUTIONS

The research project was designed by A. H. F., M. C., and T. B. S. Analyses were conducted by A. H. F. with input from M.C. and T. B. S. A. H. F. wrote the paper with T. B. S. All authors read and approved the final manuscript.

