## Supplemental methods, results, tables ane figures for "Error, noise and bias in *de novo* transcriptome assemblies"

**This PDF file includes:**

Supplementary Methods S1 – S5
Supplementary Results S1 – S3
Supplementary Tables S1 – S8
Supplementary Figures S1 – S19

### Methods

#### Methods S1: Short read processing

After an initial assessment of sequences reads with Fastqc (http://www.bioinformatics.babraham.ac.uk/projects/fastqc/), we performed kmer-based error correction with Rcorrector (1), followed by adapter and quality trimming with TrimGalore (https://www.bioinformatics.babraham.ac.uk/projects/trim_galore/), setting both the minimum phred quality score and adapter filtering stringency parameter to 5, and the minimum post-trimming read length to 36. We based our quality trimming threshold upon previous findings that performing stricter quality filtering can negatively impact assemblies (2). A second run of Fastqc was used to confirm that there was no remaining adapter sequence in the filtered reads.

#### Methods S2: Evaluation of assembly quality

To perform an initial assessment of assembly quality, we calculated standard metrics typically reported in de novo transcriptome studies. First, we quantified the total number of contigs ("transcripts" in Trinity nomenclature), the number of Trinity-defined "genes" comprising contigs deriving from the same graph, $N_{50}$, and the percent of contigs that were successfully mapped to the genome. We then assessed read support for the assembly, by mapping reads back to the assembly to assess the proportion that mapped concordantly, using methods specified by the developers of Trinity (https://github.com/trinityrnaseq/trinityrnaseq/wiki/RNA-Seq-Read-Representation-by-Trinity-Assembly). Finally, we quantified the number of completely covered, fragmented, and missing single copy conserved orthologs using the program BUSCO v. 3.0.2 (3), using the vertebrate ortholog model set for all assemblies but the fly, for which we used the arthropod set.

#### Methods S3: Reference genomes

For the purpose of benchmarking de novo transcriptome assemblies, and comparing assembly and reference-based expression estimates, we downloaded from ENSEMBL genome and gtf annotation files for house mouse, *Mus musculus* C57BL/6J (GRCm38).

#### Methods S4: Assembly and read functional composition

We first mapped contigs to their respective reference genomes with GMAP v. 2016-06-30 (4). We then used Bedtools v. 2.25 (5) to sort, and merge mapped intervals such that each reported interval consisted of unique genomic nucleotides. Mapped intervals for GMAP alignments were converted to bed format with the bamToBed function, with the '-splitD' argument such that each mapped interval for a contig—excluding alignment gaps corresponding to putative splicing events—was converted into a separate row.

To extraction functional information at the nucleotide level, we converted gtf annotation files to bed format using BEDOPS v.2.4.19 (6) using the gtf2bed command. Unique genomic bases were then classified as follows.

CDS—We extracted intervals for CDS entries for transcripts that belonged to the "protein_coding" transcript biotype.

UTR—To obtain UTR intervals, we first created a bed file representing the union of all 3' UTR, 5' UTR, CDS, start codon, and start codon intervals from protein coding transcripts. We then created a separate union of CDS, start codon, and stop codon. Finally, we subtracted the latter from the former to obtain UTR intervals. This approach prioritizes classification to CDS and stop/start codons.

Start/stop codons—Generated in the same fashion as for CDS.

Noncoding elements—We first extracted all intervals for elements corresponding to transcripts that did not have "protein_coding" as their biotype, excluding "gene" and "transcript" level entries (which correspond to the boundaries of elements, and thus are inclusive of noncoding transcript introns). We then used BEDTOOLS to subtract from this unfiltered interval set all intervals corresponding to UTR, CDS, and start/stop codons of protein coding sequence. In this way, we prioritize assignment to elements from protein coding transcripts over non-coding sequences.

Introns—To obtain putative intronic sequence, we take the transcript-level entries for protein coding transcripts (representing the outer boundaries of those transcripts) and with Bedtools subtract from them the union of all non-intron elements from both protein coding and noncoding transcripts. This prioritizes classification of nucleotides as belonging to functional elements in a genome, whether protein coding or noncoding, over intronic bases.

Noncoding introns—Noncoding sequences include elements with exon/intron structure such as spliced pseudogenes. To identify these sequences, we subtract from the transcript boundaries of all non-coding transcripts the intervals for protein coding transcript functional elements (CDS, UTR, start/stop codon), their introns, as well as the functional elements of non-coding transcripts.

Unannotated/intergenic—These genomic intervals correspond to two classes of bases: 1) those that fall entirely outside of annotation boundaries, and thus are either true intergenic sequences or originate from un-annotated functional elements, and 2) those that fall within gene boundaries of annotations, but overlap neither with exons nor introns, i.e. they occur in intervals between the stop of one transcript and the start of another that are not covered by any other transcript sequence or intron. We identify the former as those proportions of GMAP alignments of contigs falling outside of annotation boundaries. We identify the latter by subtracting the union of all functional elements and introns as defined above from bed file derived from the original annotation, which includes gene-level annotation entries.

For individual assemblies, we classified bases as belonging to particular types of elements by using the Bedtools intersectBed function to intersect the merged GMAP alignment intervals with the bed files corresponding to each of the functional classes defined above. To obtain intergenic/unannotated intervals, we obtained those falling outside of annotation boundaries by subtracting the full annotation bed file from the merged GMAP alignment bed file. Intergenic intervals falling within the annotation boundaries were defined by intersecting the alignment bed file with that for the intergenic intervals falling within the reference annotation as defined above. We first calculated assembly composition by considering all mappings of assembly contigs. To confirm that findings were not biased by a small number of contigs that mapped to many locations in the genome, we also repeated this analysis excluding all multi-mapping contigs.

To assess the functional composition of sequence reads, we randomly sub-sampled one million paired-end reads, and aligned them to their respective genomes using HISAT2 v. 2.0.4 (7). We then performed the same hierarchical classification as was done for assembled contigs, calculating frequencies for individual sequence fragments, i.e. across both sequence reads when both were properly mapped. For multi-mapped reads, we calculated a weighted average across all mappings. Visualization of functional composition was obtained by randomly sub-sampling 100,000 reads that successfully aligned to the genome.

**Methods S5: Genotyping**

For the *Mus* samples representing pools, following the GATK developers' recommendations, we also used the --sample_poidy argument, with a value set to two times the number of individuals in the pool. We edited the variant calling script to include the --sample_ploidy argument. We did not use either base quality score or variant quality score recalibration, to reflect the fact that the truth sets needed for these methods are rarely, if ever, available for ecological genomics data. We applied the same strand bias, depth, and quality-by-depth filters to raw variant calls from both methods to produce a final filtered set. To define callable sites for the map-to-reference approach, we extracted read alignments to annotated exonic regions, and then calculated the number of exonic sites with a coverage depth ≥ 5. For SuperTranscripts, we performed similar operations, defining exonic coordinates from mappings of SuperTranscripts to the *Mus* genome using GMAP, and callable sites within exons in the same way as with map-to-reference, i.e. coverage depth ≥ 5. We defined heterozygous calls in our filtered vcf files as all bi-allelic SNP sites that passed filtering criteria. In cases where sample ploidy was greater than diploid, we collapsed genotypes into the union of all unique recorded alleles.

We obtained exonic regions for SuperTranscripts by mapping them to the *Mus* genome using GMAP, then intersected these mappings with a bed file of exonic intervals with Bedtools. We used a slightly newer version of GMAP (v. 2018-07-04**)** than we did for the contig composition, which fixed minor issues with CIGAR string handling that had the potential to generate spurious genotype errors. Python scripts (https://github.com/harvardinformatics/TranscriptomeAssemblyEvaluation) were then used make transformations between genomic and SuperTranscript coordinates, for the purpose of extracting SuperTranscript genotypes and writing genotype bed files back to genomic coordinates.

Because we ran GATK such that only called genotypes—those containing an allele other than the genome or SuperTranscript reference sequence—there can be instances where the underlying map-to-reference and SuperTranscript genotypes are concordant but comparison of called genotypes will incorrectly suggest a genotyping error. For example, if a sample is fixed for an alternative allele relative to the reference genome, in the map-to-reference approach a fixed-alternative genotype will be called. However, if the SuperTranscript assembly contains this alternative allele in its sequence, no genotype would be called when mapping to SuperTranscripts. Conversely, if a sample is fixed for the allele observed in the reference genome, and there is an assembly error in a SuperTranscript such that a different, albeit incorrect, nucleotide is present at the homologous position, no genotype will be called with map-to-reference, but a fixed-alternative genotype will be called relative to a SuperTranscript. To prevent these types of events from being called errors, we updated the sets of observed alleles for both approaches by 1) updating the map-tor-reference allele to the observed nucleotide in the reference sequence if no genotype was called and, similarly, 2) when no SuperTranscript genotype is called, updating the SuperTranscript alleles to those observed in SuperTranscripts that align to the genomic position in question. In the latter case, if no SuperTranscript alignment overlaps a map-to-reference polymorphic site, no allele updating can take place.

In order to determine if SuperTranscripts with either high false negative, false positive or overall genotyping error rates were enriched for particular functions, we performed statistical enrichment tests for Biological Process, Molecular Function and Cellular Component Gene Ontology categories using Panther (8). To carry out these tests we used the Bedtools intersectBed fuction (with the -wo flag) to intersect bed-formatted GMAP alignments of SuperTranscripts with Ensembl transcript boundaries. For each SuperTranscript, we then defined the "best hit" from this intersection as the annotated transcripts with the largest number of bases overlapping the SuperTranscript in the alignment. The gene to which the annotated transcript belonged is then called the best-hit gene. We then input to Panther a table where the first column of each row is the Ensembl gene name, and the second column is the mean error rate for SuperTranscripts whose best hit is that Ensembl gene. Test results are reported in Table S6.

### Results

#### Results S1: Assembly quality metrics

96 – 98% of read pairs aligned to their respective assemblies, and upwards of 95% of these alignments occurred in a concordant fashion, with both reads aligning to the same contig (Table S3). Between 77 and 90% of BUSCOs were recovered in either complete or fragmented form (Table S4).

#### Results S2: Redundancy and intron retention

Considering the variation in CDS and intron content among contigs that cover a single-isoform gene, while there are plenty of cases with variable assembly of CDS among contigs, the event of highest frequency across all assemblers appears to be one where there is no variation in CDS among contigs but variable intron retention, due to either transcription of intron sequence or the inclusion of un-spliced pre-mRNA in sequencing libraries (Fig. S8). Looking across all contigs, regardless of their genomic position of origin, all assemblies contain large numbers of contigs that are nearly identical (Figs. S4 and 5). For Trinity assemblies that putatively report "genes" as components, while the redundancy is greater for contigs assembled from the same graph, there are a substantial number of contigs with high similarity that originate from different Trinity "genes" (Fig. S6). From different component graphs, Trinity will erroneously assemble haplotypes that differ only by a few SNVs (Fig. S4 B).

#### Results S3: Kallisto expression estimates

Adjusting for effective length had little effect on Kallisto CPM, and in some cases reduced the correlation between methods. Kallisto may be doing a better job at estimating effective length on transcriptome assemblies, such that its inferior performance relative to RSEM must be due to other reasons.

**Table S1**. SRA accession number and paired-end sequencing library attributes for samples used in this study.

| Accession | Species/strain | Tissue | Sampling | Read length | Platform | Stranded | Pre-filtered[a] | Reads (millions) | Citation |
|---|---|---|---|---|---|---|---|---|---|
| SRR203276 | *Mus musculus* – C57BL/6J | dendritic cells | 1 individual | 75 | GAII | RF | N | 52.6 | 1 |
| SRR2040596-7 | *M. musculus* – BALB/c | brain | Pool: 3M, 3F | 100 | HiSeq 2000 | FR | Y | 148.6 | 2 |
| ERR1101637 | *M. m. domesticus*, Massif Central, France | brain | Pool of 8 | 100 | HiSeq 2000 | None | Y | 58.6 | 3 |
| ERR1101633 | *M. m. domesticus*, Cologne/Bonn, Germany | brain | Pool of 8 | 100 | HiSeq 2000 | None | Y | 65.1 | 3 |
| ERR1101629 | *M. m. domesticus*, Ahvaz, Iran | Brain | Pool of 8 | 100 | HiSeq 2000 | None | Y | 47.5 | 3 |
| ERR1101645 | *M. m. musculus*, Almaty, Kazhakstan | brain | Pool of 8 | 100 | HiSeq 2000 | None | Y | 66.5 | 3 |

[a] Y indicates some fraction of raw reads obtained directly from SRA have lengths less that the identified sequencing strategy.

### Data Set References


1. Grabherr, M. G., *et al.* (2011) Full-length transcriptome assembly from RNA-Seq data without a reference genome. *Nature Biotechnology*, *29*, 644–652.
2. Ruiz-Orera, J., *et al*. (2015). Origins of De Novo Genes in Human and Chimpanzee. *PLoS Genetics, 11*, e1005721.
3. Harr, B., *et al.* (2016). Genomic resources for wild populations of the house mouse, *Mus musculus* and its close relative *Mus spretus*. *Scientific Data*, *3*, 160075.

**Table S2.** Summary statistics for de novo transcriptome assemblies.

| **Assembly** | **Assembled bases** | **Number contig clusters**[a] | **Number contigs** | **Median contig length** | **N50** |
|---|---|---|---|---|---|
| *Mus* dendritic cell | | | | | |
| Trinity | 78836818 | 61620 | 70375 | 440 | 2547 |
| Shannon | 160747370 | — | 106361 | 811 | 2910 |
| BinPacker | 72298327 | — | 40206 | **1174** | **3008** |
| *Mus* BALB/c brain 3M + 3F pool | | | | | |
| Trinity | 280791751 | 352295 | 457900 | 338 | 870 |
| Shannon | 682464577 | — | 582133 | **604** | **2214** |
| BinPacker | 200774995 | — | 234605 | 421 | 1611 |
| *Mus* wild outcrossed pool – Massif Central | | | | | |
| Trinity | 307912010 | 301162 | 344217 | 389 | 2028 |
| Shannon | 701015270 | — | 538212 | 565 | 2818 |
| BinPacker | 268759897 | — | 184809 | **706** | **2945** |
| *Mus* wild outcrossed pool – Iran | | | | | |
| Trinity | 287011312 | 275027 | 318316 | 389 | 2057 |
| Shannon | 640561962 | — | 492774 | 565 | 2800 |
| BinPacker | 253391372 | — | 168035 | **738** | **3021** |
| *Mus* wild outcrossed pool – Kazakhstan | | | | | |
| Trinity | 315066017 | 319029 | 363938 | 386 | 1870 |
| Shannon | 733510085 | — | 556436 | 572 | 2858 |
| BinPacker | 281584028 | — | 199379 | **675** | **2918** |
| *Mus* wild outcrossed pool – Germany | | | | | |
| Trinity | 325601291 | 314817 | 359664 | 391 | 2065 |
| Shannon | 758993487 | — | 578403 | 577 | 2804 |
| BinPacker | 286710559 | — | 195407 | **711** | **2993** |

[a] Shannon and BinPacker do not group contigs into such clusters, while Trinity labels them as putative genes.

**Table S3**. Read representation for de novo transcriptome assemblies, based upon percentages of properly paired aligned reads.

| Assembly | % properly paired | % left only | % right only | % Improperly paired | Aligned fragments | % reads aligned | % contigs aligned to genome |
|---|---|---|---|---|---|---|---|
| *Mus* dendritic cell | | | | | | | |
| Trinity | 96.68 | 0.73 | 0.46 | 2.13 | 41628192 | 94.79 | 98.6 |
| Shannon | 96.4 | 0.6 | 0.4 | 2.6 | 38619227 | 87.93 | 98.0 |
| BinPacker | 96.01 | 1.05 | 0.48 | 2.46 | 41229255 | 93.88 | 98.4 |
| *Mus* BALB/c brain pool (n=6) | | | | | | | |
| Trinity | 97.25 | 0.20 | 0.04 | 2.51 | 102809183 | 98.70 | 97.7 |
| Shannon | 95.95 | 0.16 | 0.04 | 3.85 | 103026767 | 98.90 | 97.3 |
| BinPacker | 96.65 | 0.34 | 0.05 | 2.97 | 101847596 | 97.78 | 97.4 |
| *Mus* wild outcrossed pool – FRA (n=8) | | | | | | | |
| Trinity | 97.48 | 0.46 | 0.19 | 1.87 | 57073082 | 98.16 | 98.1 |
| Shannon | 96.77 | 0.39 | 0.17 | 2.67 | 57225138 | 98.42 | 96.9 |
| BinPacker | 96.08 | 1.02 | 0.27 | 2.63 | 54972706 | 94.55 | 97.5 |
| *Mus* wild outcrossed pool – IRN[a] (n=8) | | | | | | | |
| Trinity | 97.10 | 0.52 | 0.23 | 2.15 | 45988313 | 97.84 | 98.1 |
| Shannon | 96.45 | 0.43 | 0.21 | 2.91 | 46174692 | 98.23 | 97.0 |
| BinPacker | 95.97 | 1.00 | 0.29 | 2.73 | 44201170 | 94.04 | 97.6 |
| *Mus* wild outcrossed pool – KZK[a] (n=8) | | | | | | | |
| Trinity | 98.05 | 0.34 | 0.13 | 1.47 | 64844818 | 98.20 | 97.6 |
| Shannon | 97.57 | 0.28 | 0.12 | 2.03 | 65072575 | 98.55 | 96.7 |
| BinPacker | 97.01 | 0.83 | 0.20 | 1.96 | 62426676 | 94.54 | 97.2 |
| *Mus* wild outcrossed pool – DEU[a] (n=8) | | | | | | | |
| Trinity | 97.21 | 0.49 | 0.20 | 2.11 | 63360450 | 98.16 | 98.2 |
| Shannon | 96.51 | 0.40 | 0.18 | 2.91 | 63578432 | 98.50 | 96.9 |
| BinPacker | 95.94 | 0.98 | 0.29 | 2.79 | 61266479 | 94.92 | 97.6 |

[a] Assembled only with Trinity

**Table S4.** Coverage of core genes (BUSCOs) by de novo transcriptome assemblies. Assemblies were searched for 2586 vertebrate BUSCOs.

| Assembly | Complete single copy | Complete duplicated | Complete total | % Recovered | Fragmented | Missing |
|---|---|---|---|---|---|---|
| Mus dendritic cell[a] | | | | | | |
| Trinity | 1452 | 648 | 2100 | 81.2 | 125 | 361 |
| Shannon | 873 | 1242 | 2115 | 81.8 | 107 | 364 |
| BinPacker | 1562 | 574 | 2136 | 82.6 | 88 | 362 |
| *Mus* BALB/c brain pool (n=6) | | | | | | |
| Trinity | 1339 | 877 | 2216 | 85.7 | 244 | 126 |
| Shannon | 334 | 1892 | 2226 | 86.1 | 234 | 126 |
| BinPacker | 1244 | 749 | 1993 | 77.1 | 444 | 149 |
| *Mus* wild outcrossed pool – Massif Central (n=8) | | | | | | |
| Trinity | 1294 | 1044 | 2338 | 90.4 | 146 | 102 |
| Shannon | 540 | 1770 | 2310 | 89.3 | 177 | 99 |
| BinPacker | 1184 | 1060 | 2244 | 86.8 | 201 | 141 |
| *Mus* wild outcrossed pool – Iran[a] (n=8) | | | | | | |
| Trinity | 1253 | 1044 | 2297 | 88.8 | 170 | 119 |
| Shannon | 525 | 1773 | 2298 | 88.9 | 173 | 115 |
| BinPacker | 1138 | 1127 | 2265 | 87.6 | 174 | 147 |
| *Mus* wild outcrossed pool – Kazakhstan[a] (n=8) | | | | | | |
| Trinity | 1295 | 1055 | 2350 | 90.1 | 145 | 91 |
| Shannon | 488 | 1852 | 2340 | 90.4 | 156 | 90 |
| BinPacker | 1146 | 1134 | 2280 | 88.2 | 172 | 134 |
| *Mus* wild outcrossed pool – Germany[a] (n=8) | | | | | | |
| Trinity | 1288 | 1046 | 2334 | 90.2 | 149 | 103 |
| Shannon | 536 | 1790 | 2326 | 89.9 | 155 | 105 |
| BinPacker | 1157 | 1129 | 2286 | 88.4 | 169 | 131 |

**Table S5**. For genomic positions with either a map-to-reference or associated SuperTranscript genotype call, counts and rates of genotyping errors per SuperTranscript.

| Sample | Mean # errors | Median # errors | Max # errors | Mean error frequency | Median error frequency | Max error frequency |
|---|---|---|---|---|---|---|
| MDC | 2.72 | 1 | 125 | 0.0020 | 0.0006 | 0.0497 |
| BALB/c | 2.55 | 2 | 56 | 0.0038 | 0.0014 | 0.1330 |
| FRA | 2.92 | 2 | 125 | 0.0023 | 0.0009 | 0.0980 |
| DEU | 3.02 | 2 | 82 | 0.0022 | 0.0009 | 0.1119 |
| IRN | 2.95 | 2 | 187 | 0.0022 | 0.0010 | 0.1680 |
| KZK | 2.99 | 2 | 146 | 0.0023 | 0.0010 | 0.1294 |

**Table S6.** Statistical enrichments for Gene Ontology terms for SuperTranscripts with high false positive or overall error rate. We report terms with FDR≤ 0.05 that represent the highest significant term within a hierarchy, i.e. the most general term of a hierarchical set.Terms with negative enrichments—deficits in terms in SuperTranscripts with high error rates—are not reported. Only terms with FDR≤ 0.05 are reported.

| Sample | GO term | # Genes | FDR |
|---|---|---|---|
| **FRA** | | | |
| **FP** | **Biological Process** | | |
| | regulation of synaptic structure or aactivity (0050803) | 221 | $4.34 \times 10^{-2}$ |
| | plasma membrane bounded cell projection morphogenesis (0120039) | 338 | $3.67 \times 10^{-2}$ |
| | **Cellular Component** | | |
| | glutamatergic synapse (0098978) | 411 | $7.05 \times 10^{-5}$ |
| | postsynaptic density (0014069) | 320 | $6.46 \times 10^{-4}$ |
| | ribonucleoprotein complex (1990904) | 495 | $1.62 \times 10^{-2}$ |
| | somatodendritic compartment (0036477) | 725 | $2.51 \times 10^{-2}$ |
| **Error rate** | **Biological process** | | |
| | antigen processing and presentation of endogenous peptide antigen via MHC class I viaER pathway, TAP-independent (0002486) | 10 | $1.31 \times 10^{-3}$ |
| | antigen processing and presentation of endogenous peptide antigen via MHC class Ib (0002476) | 11 | $6.70 \times 10^{-3}$ |
| | positive regulation of T cell mediated cytotoxicity (0001916) | 13 | $9.29 \times 10^{-3}$ |
| | centrosome cycle (0007098) | 34 | $1.56 \times 10^{-2}$ |
| | defense response to bacterium (0042742) | 47 | $2.42 \times 10^{-2}$ |
| | **Molecular function** | | |
| | structural constituent of ribosome (0003735) | 61 | $3.34 \times 10^{-3}$ |
| | beta-2-microglobulin binding (GO0030881) | 6 | $2.24 \times 10^{-2}$ |
| | T cell receptor binding (0042608) | 9 | $4.87 \times 10^{-2}$ |
| | **Cellular component** | | |
| | cytosolic large ribosomal subunit (0022625) | 24 | $1.10 \times 10^{-3}$ |
| | MHC class I protein complex (0042612) | 6 | $1.01 \times 10^{-2}$ |
| **DEU** | | | |
| **Error rate** | **Molecular function** | | |
| | structural constituent of ribosome (0003735) | 58 | $1.22 \times 10^{-5}$ |
| | **Cellular component** | | |
| | cytosolic large ribosomal subunit (0022625) | 22 | $3.04 \times 10^{-3}$ |
| | cytosolic small ribosomal subunit (GO?0022627) | 22 | $4.16 \times 10^{-2}$ |
| **IRN** | **Molecular function** | | |
| | structural constituent of ribosome (0003735) | 67 | $2.81 \times 10^{-6}$ |
| | GTPase activity (J0003924)) | 148 | $2.72 \times 10^{-2}$ |

| | | | |
|---|---|---|---|
| | **Cellular component** | | |
| | cytosolic large ribosomal subunit (0022625) | 31 | $1.76 \times 10^{-4}$ |
| | cytosolic small ribosomal subunit (0022627) | 21 | $1.75 \times 10^{-2}$ |
| | inner mitochondrial membrane protein complex (0098800) | 41 | $3.96 \times 10^{-2}$ |
| **KZK** | | | |
| **Error rate** | **Cellular component** | | |
| | cytosolic large ribosomal subunit (0022625) | 25 | $7.79 \times 10^{-3}$ |
| | small ribosomal subunit (0015935) | 33 | $2.61 \times 10^{-2}$ |

**Table S7**. Spearman rank correlations between map-to-reference and map-to-denovo CPM and TPM estimated with RSEM, for BLAT best hit contigs and their respective reference isoforms.

| | Best hit-to-transcript | | | Best hit-to-gene |
|---|---|---|---|---|
| **Sample** | **CPM** | **TPM** | **Rescaled CPM** | **Rescaled CPM** |
| MDC | | | | |
| Trinity | 0.259 | 0.319 | 0.336 | 0.637 |
| Shannon | 0.070 | 0.073 | 0.089 | 0.103 |
| BinPacker | 0.227 | 0.302 | 0.315 | 0.616 |
| BALB/c | | | | |
| Trinity | 0.387 | 0.385 | 0.406 | 0.657 |
| Shannon | 0.079 | 0.045 | 0.083 | 0.023 |
| BinPacker | 0.322 | 0.314 | 0.346 | 0.543 |
| Massif Central | | | | |
| Trinity | 0.363 | 0.372 | 0.401 | 0.603 |
| Shannon | 0.282 | 0.185 | 0.303 | 0.201 |
| BinPacker | 0.282 | 0.290 | 0.325 | 0.490 |
| Germany | | | | |
| Trinity | 0.362 | 0.369 | 0.402 | 0.588 |
| Shannon | 0.284 | 0.187 | 0.305 | 0.195 |
| BinPacker | 0290 | 0.294 | 0.334 | 0.482 |
| Iran | | | | |
| Trinity | 0.373 | 0.370 | 0.412 | 0.600 |
| Shannon | 0.292 | 0.192 | 0.313 | 0.210 |
| BinPacker | 0.288 | 0.293 | 0.333 | 0.494 |
| Kazakhstan | | | | |
| Trinity | 0.381 | 0.392 | 0.415 | 0.612 |
| Shannon | 0.285 | 0.189 | 0.304 | 0.206 |
| BinPacker | 0.297 | 0.301 | 0.336 | 0.489 |

**Table S8**. Correlations between map-to-reference and map-to-de novo-transcriptome expression at the gene level, for both RSEM and Kallisto, based upon CPM, TPM, and weighted-adjusted CPM. See equation in methods for details regarding the last of these. Correlations exclude expressed reference genes where there are not associated de novo assembly contigs, i.e. genes that are missing from the assembly. Correlations are calculated for entries with TPM > 0 for either map-to-reference or map-to-de novo assembly. RSEM values in parentheses are for filtered assemblies from which contigs with TPM ≤ 1 and without a BLASTP hit to Uniref90 are removed, and which are only compared for protein coding genes with a minimum transcript length ≥ 200bp. The % missing column indicates expressed protein coding genes that were not recovered by the unfiltered transcriptome assembly. The % lost-to-filtering column indicates what fraction of expressed protein coding genes that are detected in the unfiltered *de novo* transcriptome assemblies are lost due to filtering. Bold text indicates, for each assembly, the method that produces the strongest correlation with map-to-reference expression estimates.

| **Sample** | **CPM** | | **TPM** | | **Rescaled CPM** | | | |
|---|---|---|---|---|---|---|---|---|
| | **RSEM** | **kallisto** | **RSEM** | **kallisto** | **RSEM** | **kallisto** | **% missing (TPM≥1)** | **% lost-to-filtering (TPM≥1)** |
| MDC | | | | | | | | |
| Trinity | 0.853 (0.951) | 0.858 | 0.756 (0.933) | 0.675 | 0.901 **(0.957)** | 0.823 | 20.0 (2.0) | 10.6 (6.0) |
| Shannon | 0.629 (0.897) | 0.833 | 0.567 (0.901) | 0.662 | 0.680 (0.906) | 0.784 | 21.3 (2.5) | 31.5 (27.1) |
| BinPacker | 0.824 (0.940) | 0.825 | 0.796 (0.937) | 0.754 | 0.877 (0.947) | 0.783 | 25.0 (3.3) | 9.1 (6.8) |
| BALB/c | | | | | | | | |
| Trinity | 0.869 (0.934) | 0.881 | 0.818 (0.895) | 0.819 | 0.897 **(0.938)** | 0.863 | 8.6 (0.6%) | 24.5 (11.5) |
| Shannon | 0.819 (0.846) | 0.859 | 0.790 (0.848) | 0.815 | 0.854 (0.853) | 0.826 | 8.8 (0.7) | 32.5 (19.1) |
| BinPacker | 0.835 (0.877) | 0.846 | 0.801 (0.864) | 0.803 | 0.870 (0.883) | 0.824 | 12.4 (1.7) | 20.6 (10.5) |
| FRA | | | | | | | | |
| Trinity | 0.850 (0.927) | 0.861 | 0.809 (0.928) | 0.790 | 0.903 **(0.933)** | 0.841 | 11.8 (2.0) | 20.6 (10.6) |
| Shannon | 0.849 | 0.857 | 0.812 | 0.785 | 0.906 | 0.819 | 11.5 (1.6) | 24.7 (14.1) |
| BinPacker | 0.815 (0.888) | 0.821 | 0.808 (0.902) | 0.786 | 0.872 (0.896) | 0.793 | 18.3 (3.2) | 14.1 (10.5) |
| DEU | | | | | | | | |
| Trinity | 0.848 (0.925) | 0.859 | 0.804 (0.926) | 0.788 | 0.903 **(0.933)** | 0.838 | 11.7 (2.0) | 21.9 (11.5) |
| Shannon | 0.847 (0.879) | 0.855 | 0.812 (0.894) | 0.784 | 0.905 (0.888) | 0.813 | 11.3 (1.6) | 25.6 (14.9) |
| BinPacker | 0.811 (0.880) | 0.818 | 0.808 (0.900) | 0.784 | 0.873 (0.890) | 0.789 | 17.6 (3.1) | 14.7 (10.5) |
| IRN | | | | | | | | |
| Trinity | 0.853 (0.929) | 0.862 | 0.806 (0.928) | 0.785 | 0.906 **(0.937)** | 0.842 | 12.0 (2.0) | 18.9 (10.0) |
| Shannon | 0.847 (0.893) | 0.854 | 0.804 (0.902) | 0.777 | 0.904 (0.902) | 0.817 | 11.5 (1.7) | 22.1 (12.6) |
| BinPacker | 0.809 (0.888) | 0.816 | 0.799 (0.901) | 0.773 | 0.869 (0.898) | 0.787 | 19.0 (3.6) | 13.1 (10.2) |
| KZK | | | | | | | | |
| Trinity | 0.852 (0.926) | 0.862 | 0.810 (0.925) | 0.791 | 0.903 **(0.933)** | 0.841 | 10.9 (2.0) | 22.8 (12.0) |
| Shannon | 0.852 (0.885) | 0.858 | 0.814 (0.891) | 0.793 | 0.907 (0.893) | 0.824 | 10.7 (1.8) | 25.7 (14.8) |
| BinPacker | 0.816 (0.894) | 0.821 | 0.809 (0.908) | 0.788 | 0.872 (0.902) | 0.795 | 16.8 (3.4) | 14.9 (10.7) |

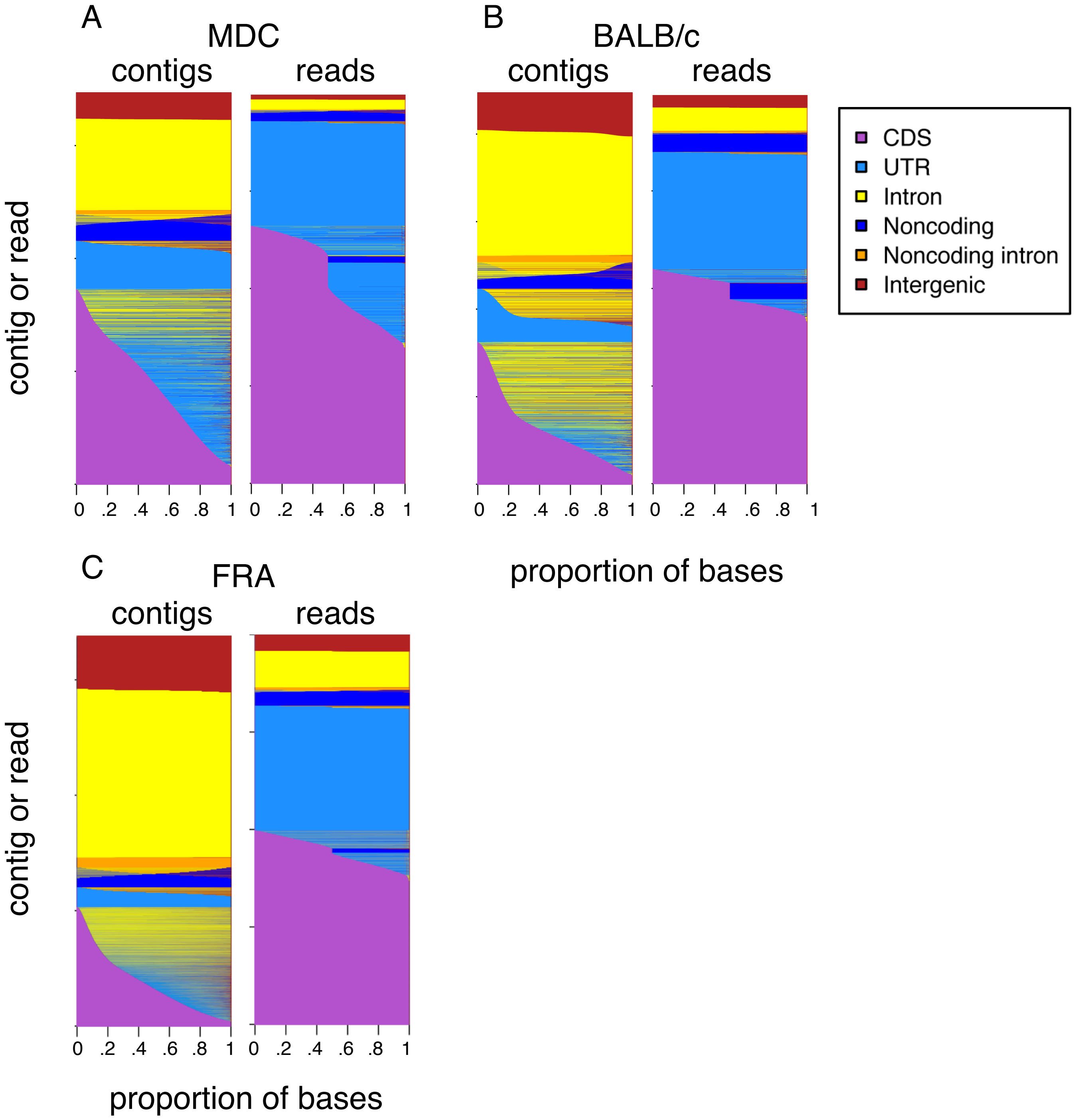


**Fig. S1.** Base composition by Trinity contig (left panels), and sequence reads (right panels) for (A) *M. musculus* dendritic cells, (B) *M. musculus* BALB/c strain brain, and (C) wild *Mus* brain pool from Massif Central, France. For each panel, each y-axis row represents a single contig or read. Sequence panels for reads are based upon 100,000 randomly sampled mapped reads. Intersections of contigs and reads with annotations are based upon mapping with GMAP and HISAT2, respectively (see Methods for details). For visualization purposes, sequence proportions are arranged from the bottom to the top of a panel, sorted in descending order, consecutively by CDS, UTR, non-coding, non-coding intron, intron, and intergenic proportions.

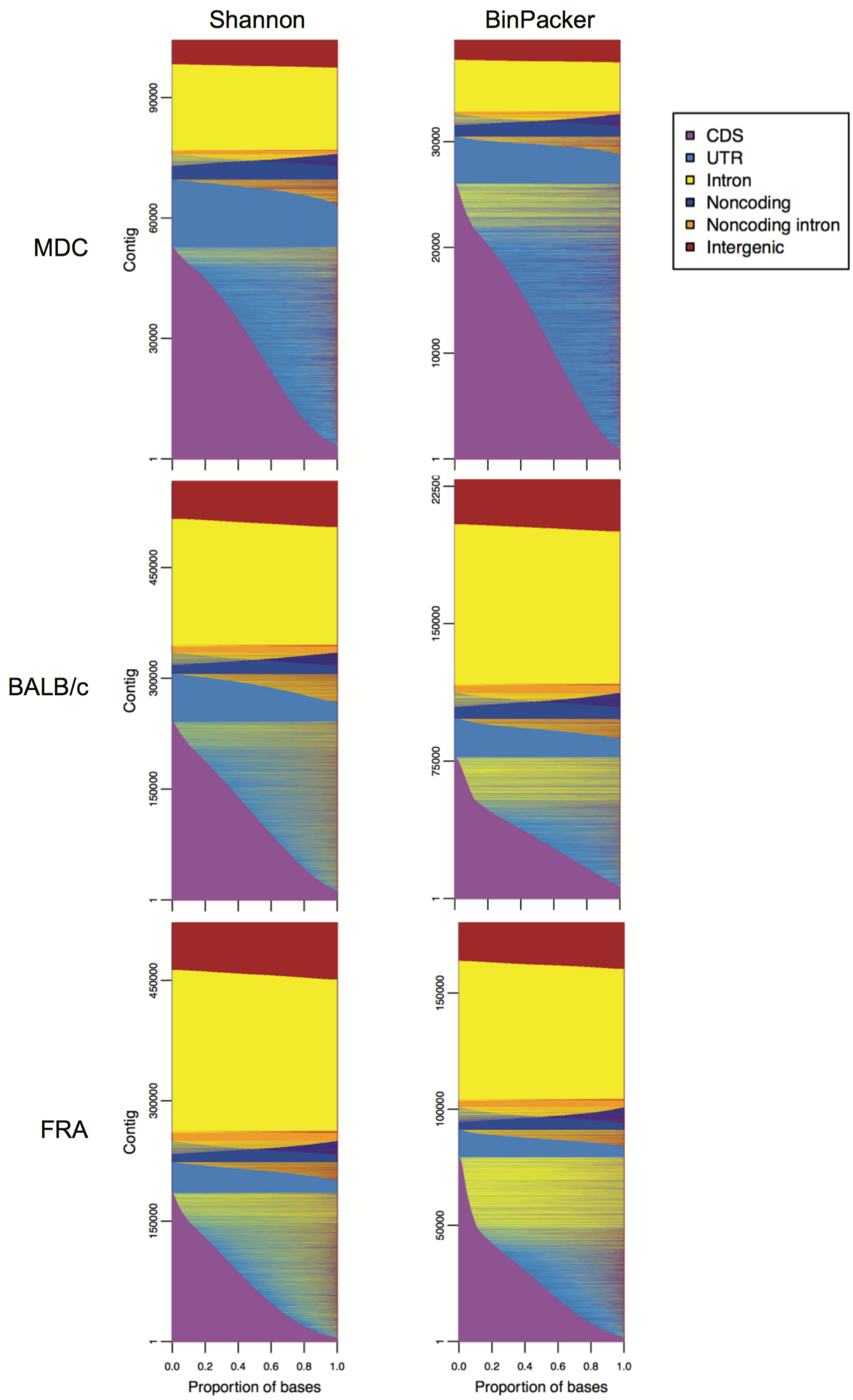


**Fig. S2.** Base composition by contig for Shannon (left panels), and BinPacker (right panels) for MDC (top row), BALB/c (middle row), and FRA (bottom row). For each panel, each y-axis row represents a single contig or read. Sequence panels for reads are based upon 100,000 randomly sampled mapped reads. Intersections of contigs and reads with annotations are based upon mapping with GMAP and HISAT2, respectively (see **Methods** for details).

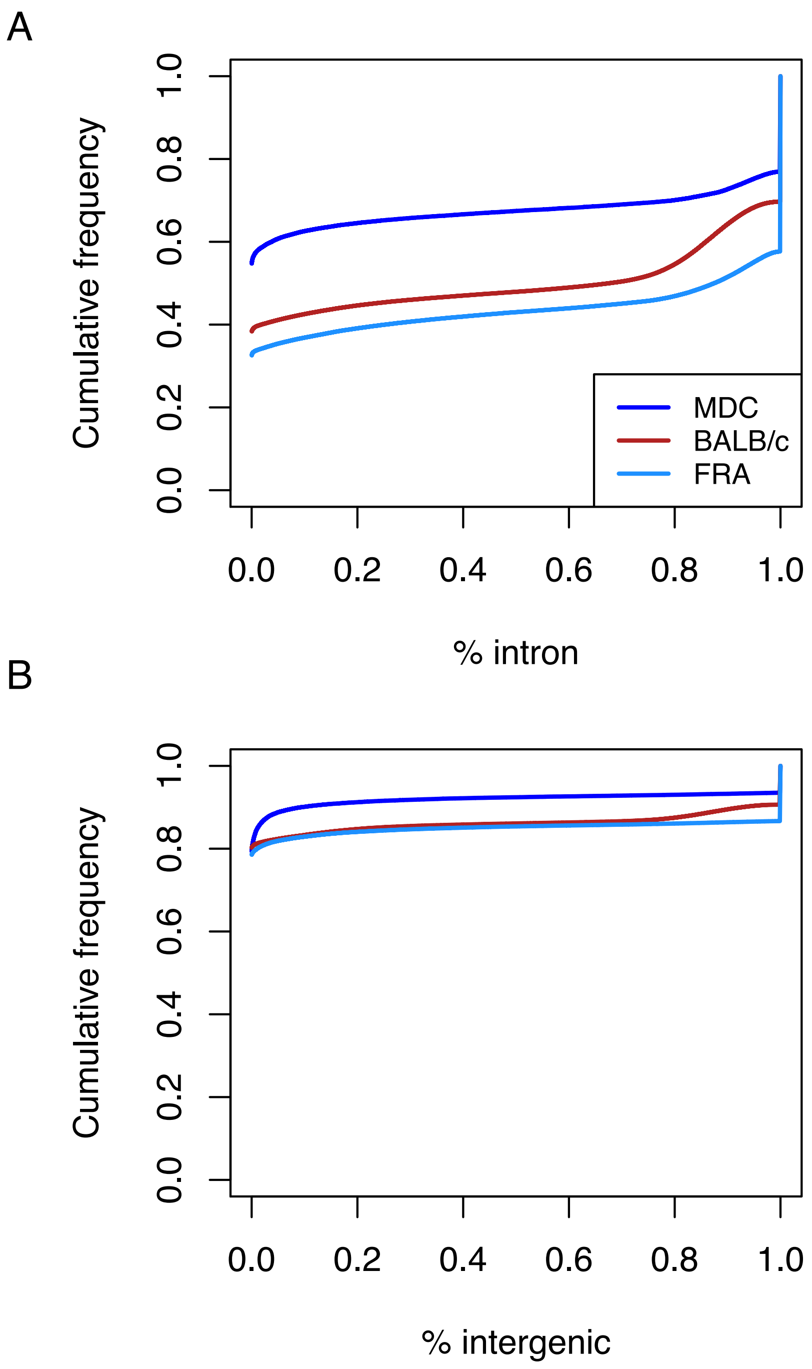


**Fig. S3.** Cumulative frequency ecdf curves of (A) intron and (B) intergenic proportions of Trinity contigs.

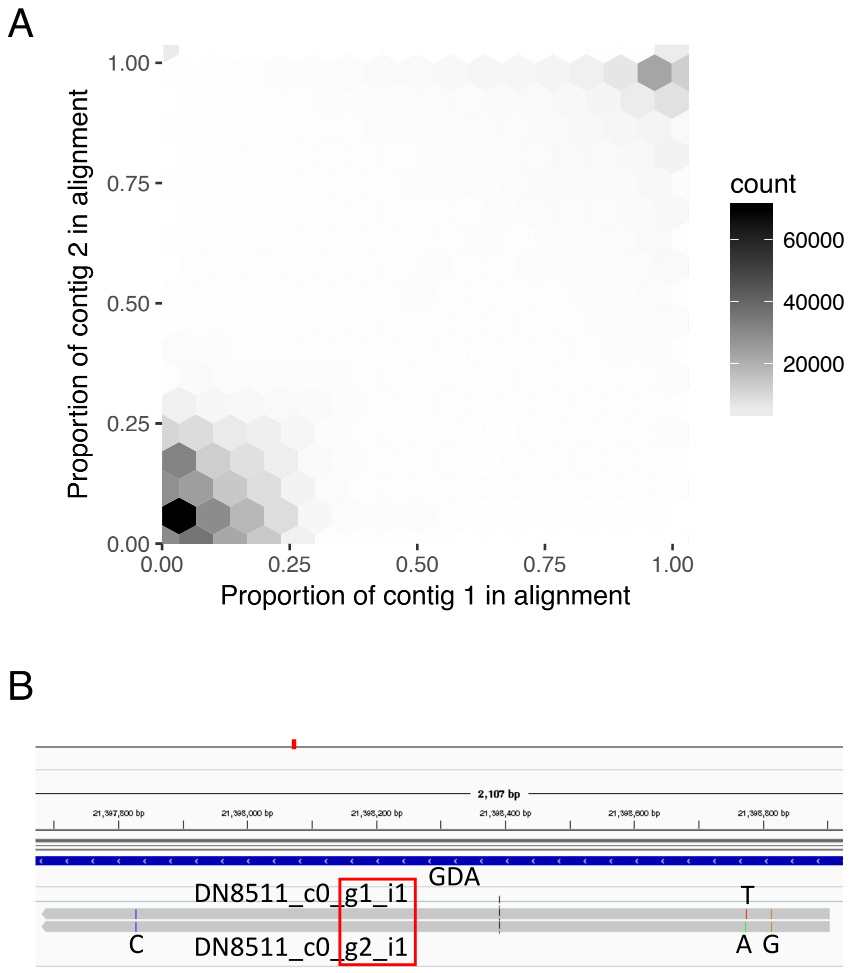


**Fig. S4.** (A) For FRA Trinity assembly, hexbin 2D histogram of the proportion of contig lengths comprising BLAT pairwise alignments resulting from all-by-all searches. (B) Example alignment to the Mus genome, to a protein-coding transcript of the GDA gene, of nearly identically sized (1220 bp) FRA Trinity contigs. These contigs have a BLAT-based 99.8 % identity in an alignment equal to the entirety of the contig length, and which are assembled from different Trinity components (i.e. “genes”), as indicated by the red box around their labels. The contigs differ by only two adjacent SNPs (A and T), and also share fixed C (far left) and G (far right) differences relative to the reference genome. Thus, different Trinity components have assembled different haplotypes.

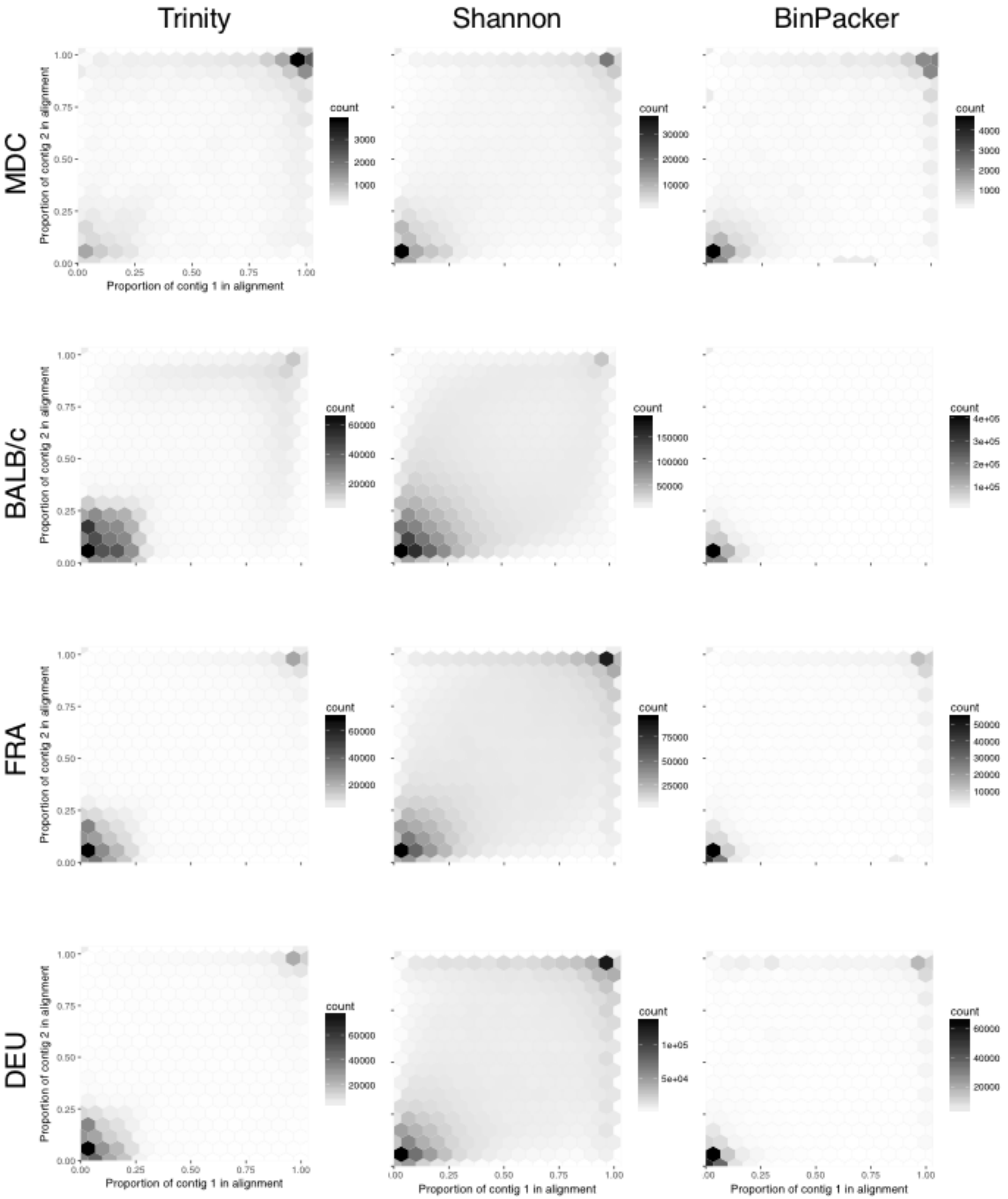

Trinity
Shannon
BinPacker
MDC
BALB/c
FRA
DEU
Proportion of contig 2 in alignment
Proportion of contig 1 in alignment
count

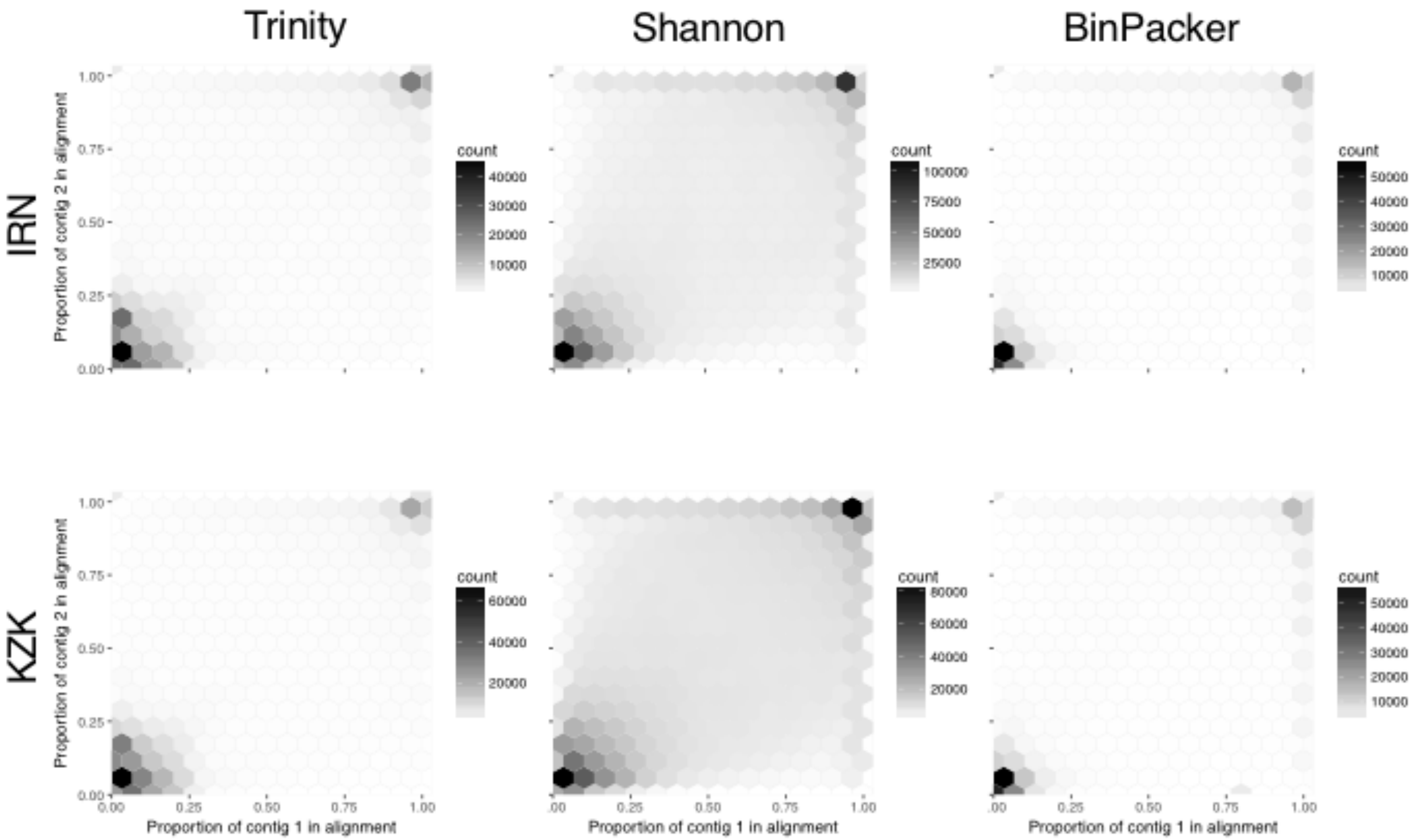


**Fig. S5.** Joint frequency hexbin plot of the proportion of query and target contig lengths comprised of alignments using a BLAT all-by-all search for all *Mus* transcriptome assemblies. Larger counts in lower left and upper right corners indicate cases where alignments represent small, and large proportions of both aligned sequences, respectively.

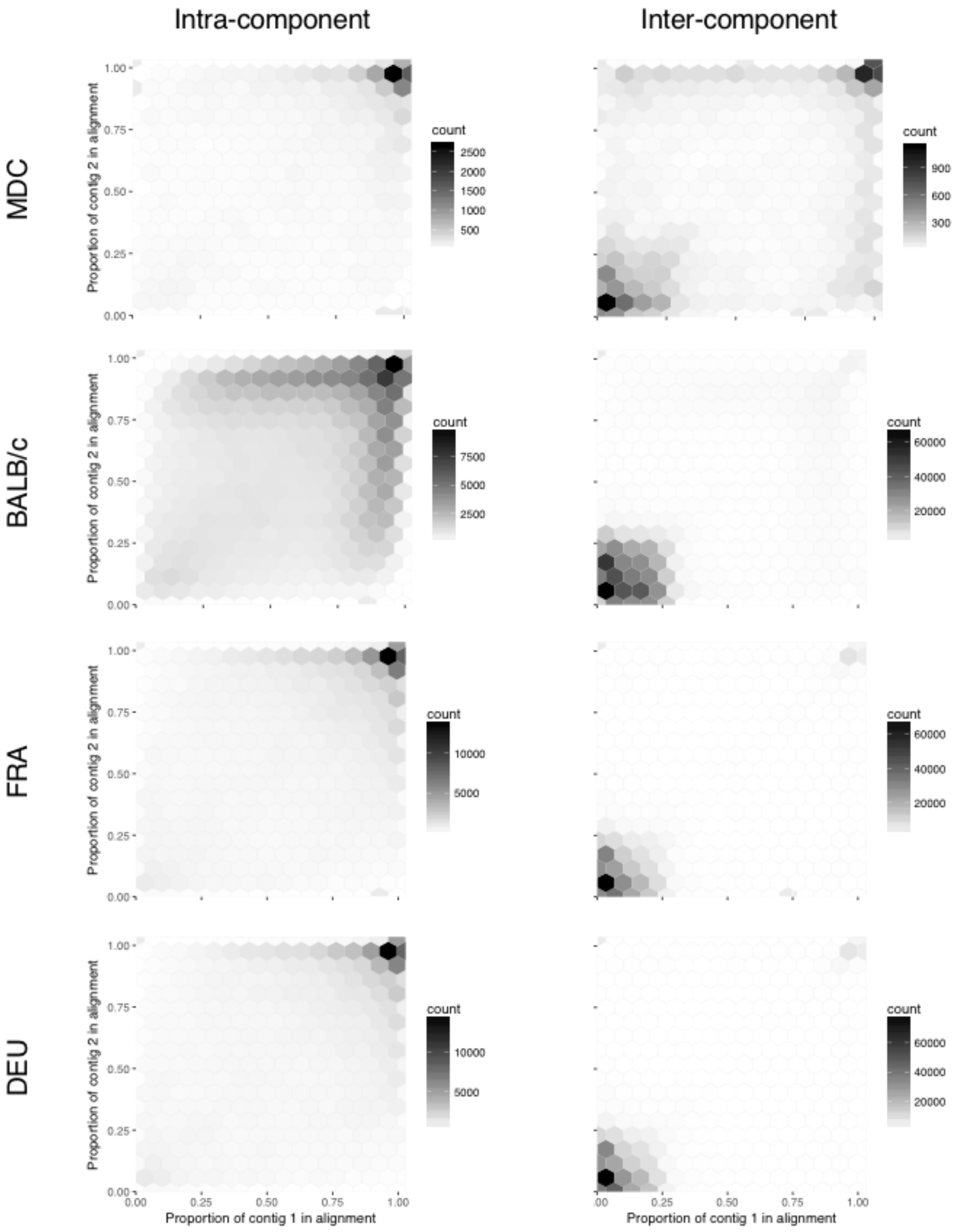
Intra-component
Inter-component
MDC
BALB/c
FRA
DEU
Proportion of contig 2 in alignment
Proportion of contig 1 in alignment
count
0.00
0.25
0.50
0.75
1.00
2500
2000
1500
1000
500
900
600
300
7500
5000
2500
60000
40000
20000
10000
5000
.00

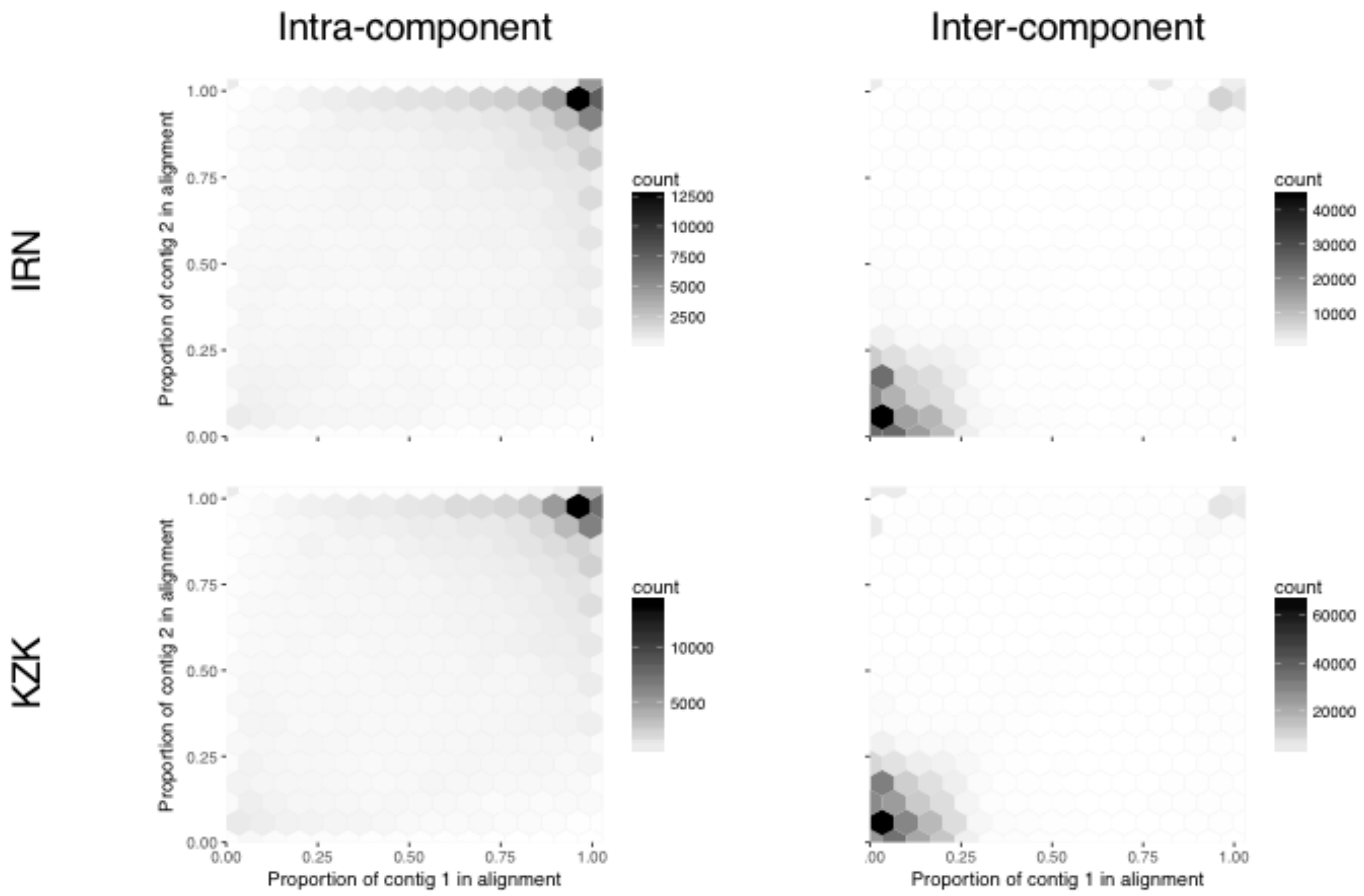


**Fig. S6.** Joint frequency hexbin plot of the proportion of query and target contig lengths comprised of alignments using a BLAT all-by-all search for *Mus* transcriptome assemblies produced with Trinity, dividing BLAT hits into those between contigs coming from the same component (i.e. Trinity "gene", left column), and different components (right column).

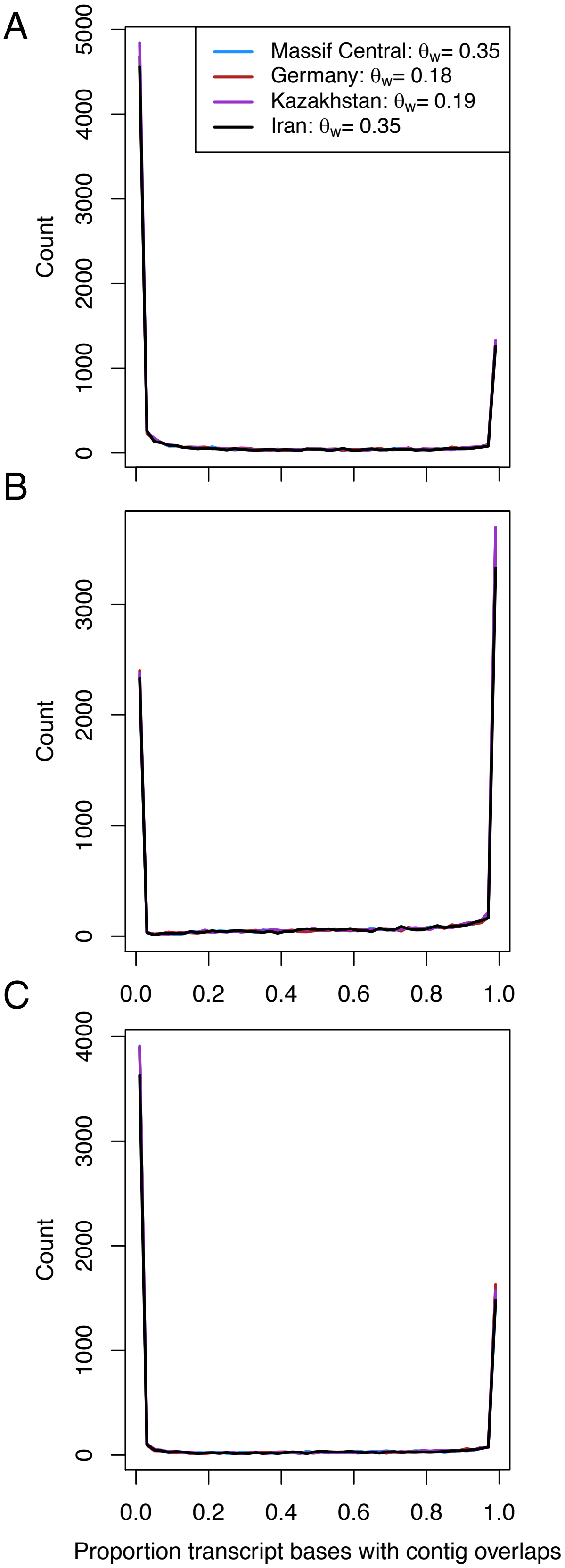


**Fig. S7.** Frequency of proportional overlaps between contigs, based upon BLAT all-by-all hits, for known Mus single isoform genes, for three 8-individual pools of wild M. m. domesticus (Iran; Massif Central, France and Germany) and one 8-individual pool of M. m. musculus (Kazakhstan), for (A) Trinity, (G) Shannon and (C) BinPacker. Estimates of Watterson's theta are those reported from whole genome sequence data in Harr et al. (2016).

A

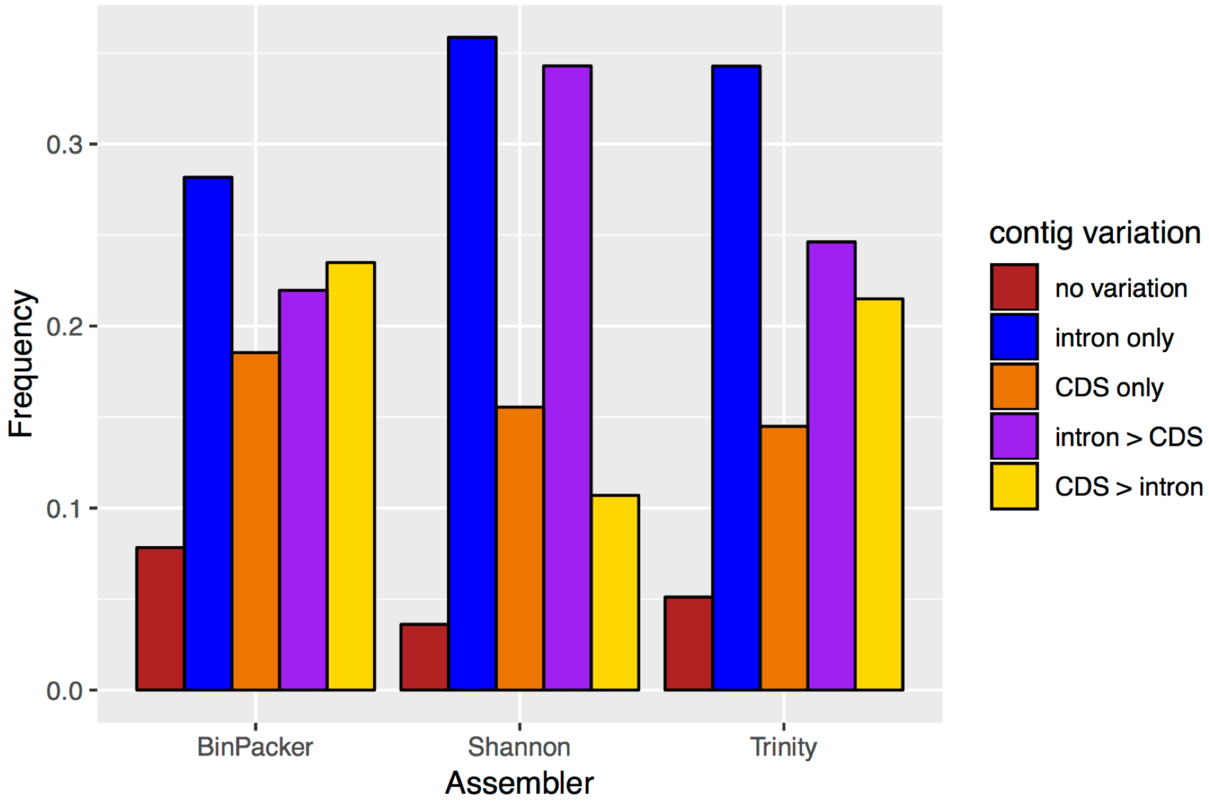


B

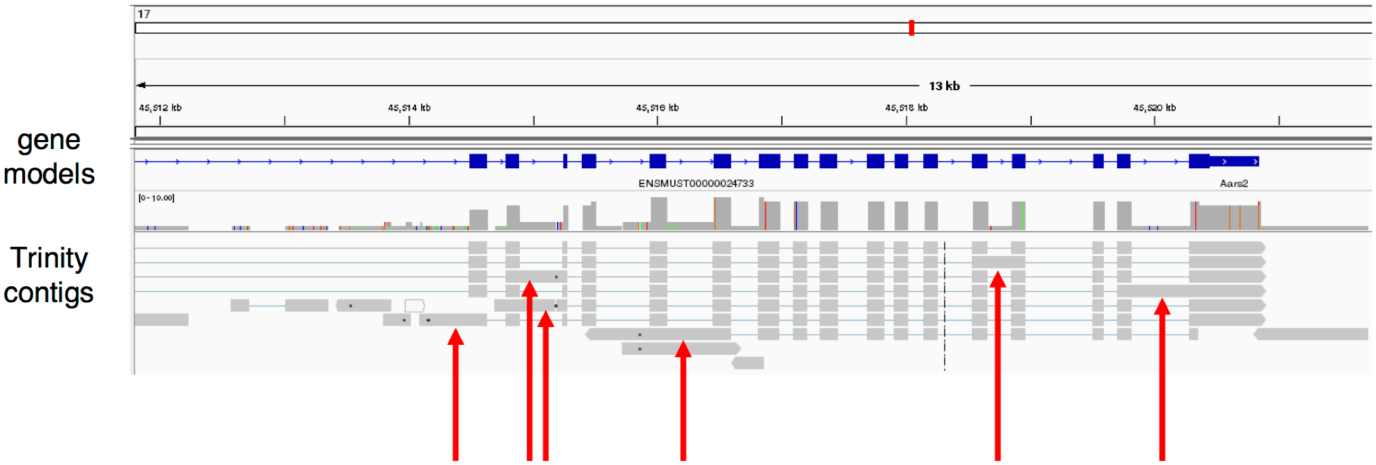


**Fig. S8**. For the FRA Trinity assembly, (A) For single-isoform genes that are covered by more than one contig at all positions, the frequency of cases where there is no variation in either intron or CDS content among contigs (no variation), no variation in CDS content but where the standard deviation of intron content among contigs is > 0 (intron only), the converse of this pattern (CDS only), cases where variation is non-zero for both classes but is greater in intron (intron > CDS), or in CDS (CDS > intron). Calculating of composition, based upon contig mapping to the genome, exclude the small number of cases where there are BLAT hits to isoform sequences but no valid GMAP alignment to the genome. (B) Genome browser view of single isoform gene AARS2 and GMAP alignments of contigs. Red arrows indicate cases where assembled contigs differ due to variation in the extent of retained intron.

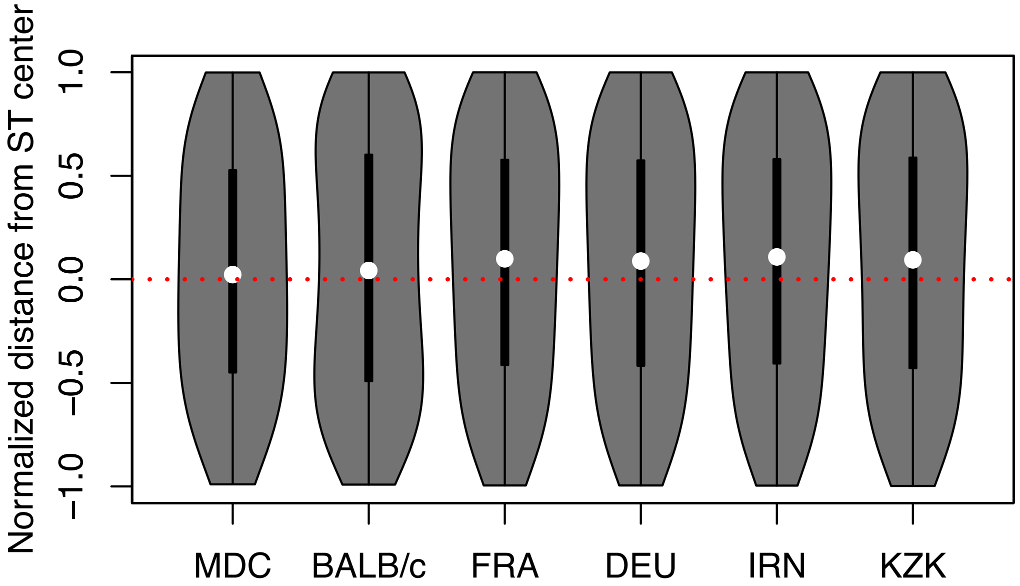


**Fig. S9.** Violin plots of the normalized (-1,1) position of SuperTranscript (ST) genotyping errors relative to the middle of an ST indicating a tendency for errors to occur at the distal end of an ST.

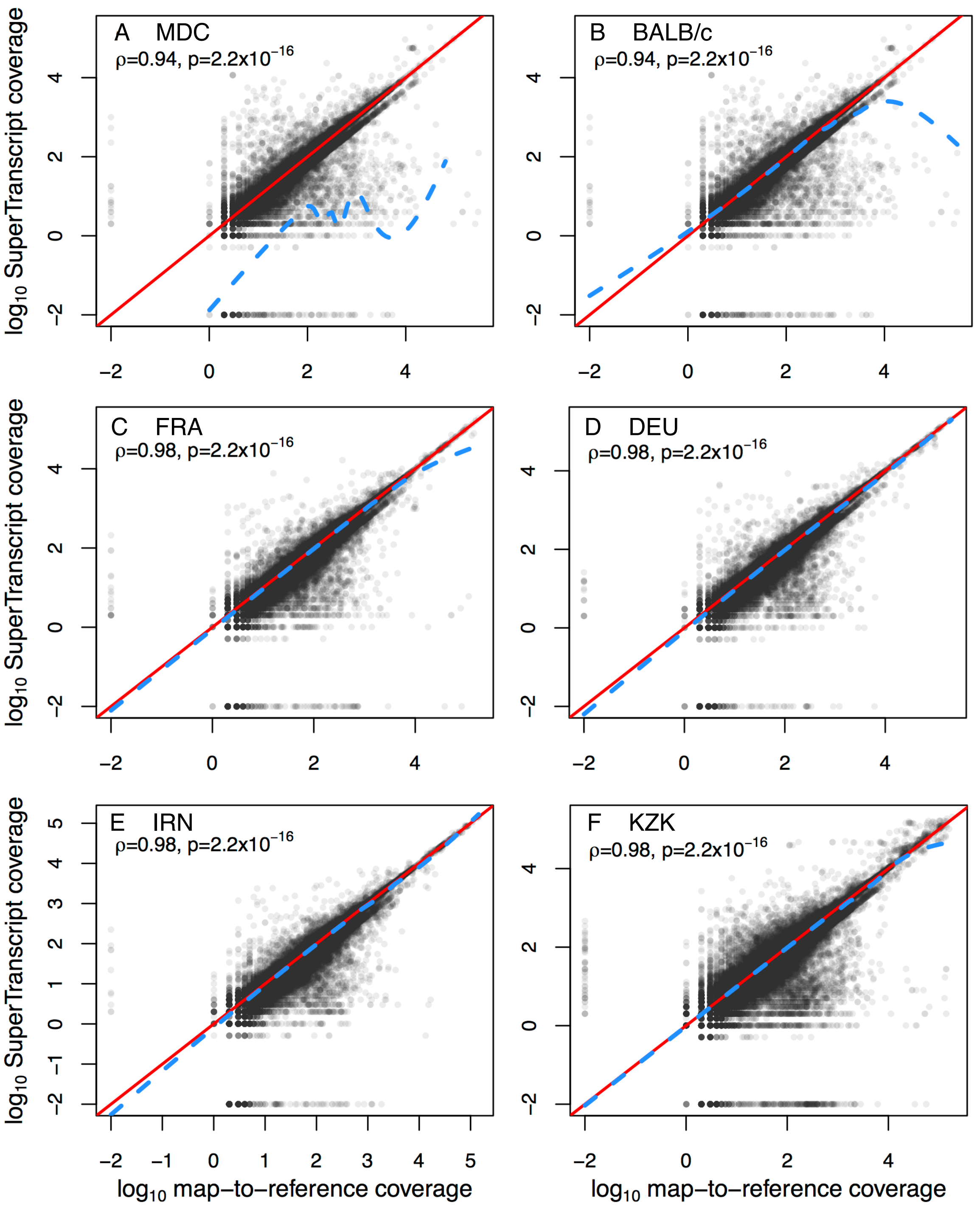


**Fig. S10.** Biplots and spearman's rank correlations of map-to-reference read depth vs. SuperTranscript read depth at positions where the genotypes called by the two methods are concordant. Red lines indicate unity, while blue lines indicated the fitted, smoothed trend. Prior to log-transformation, 0.01 is added to all values so that zero values are not omitted from the plots. Thus, log-transformed values of -2 indicate read depth of zero.

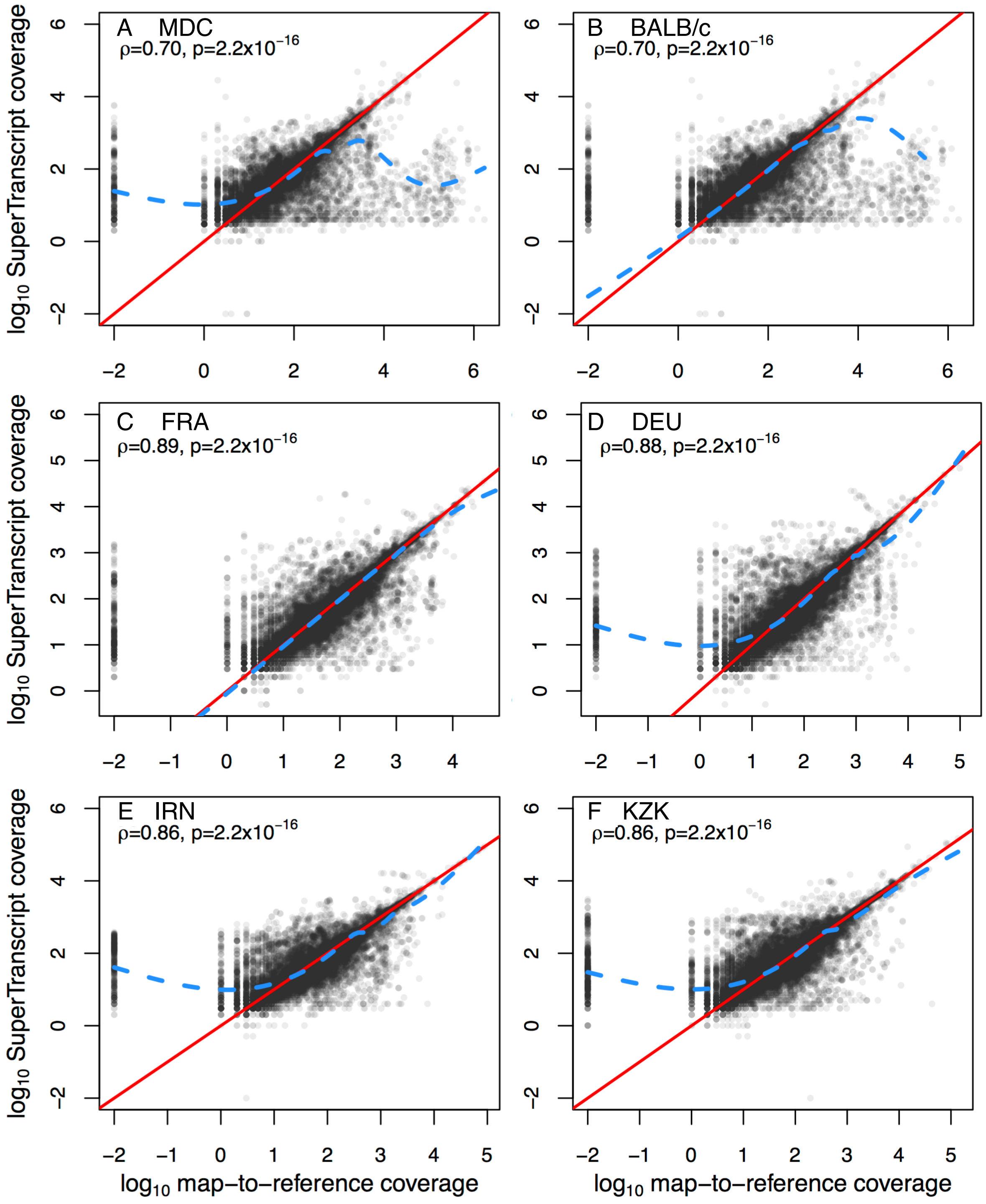


**Fig. S11.** Biplots and spearman's rank correlations of map-to-reference read depth vs. SuperTranscript read depth at positions where there are SuperTranscript (ST) false positives (FP). Blue lines indicated the fitted, smoothed trend. Prior to log-transformation, 0.01 is added to all values so that zero values are not omitted from the plots. Thus, log-transformed values of -2 indicate read depth of zero. For the BALB/c, there were two sites where ST coverage obtained from Bedtools, the coverage we used for plotting, was equal to zero, yet GATK.

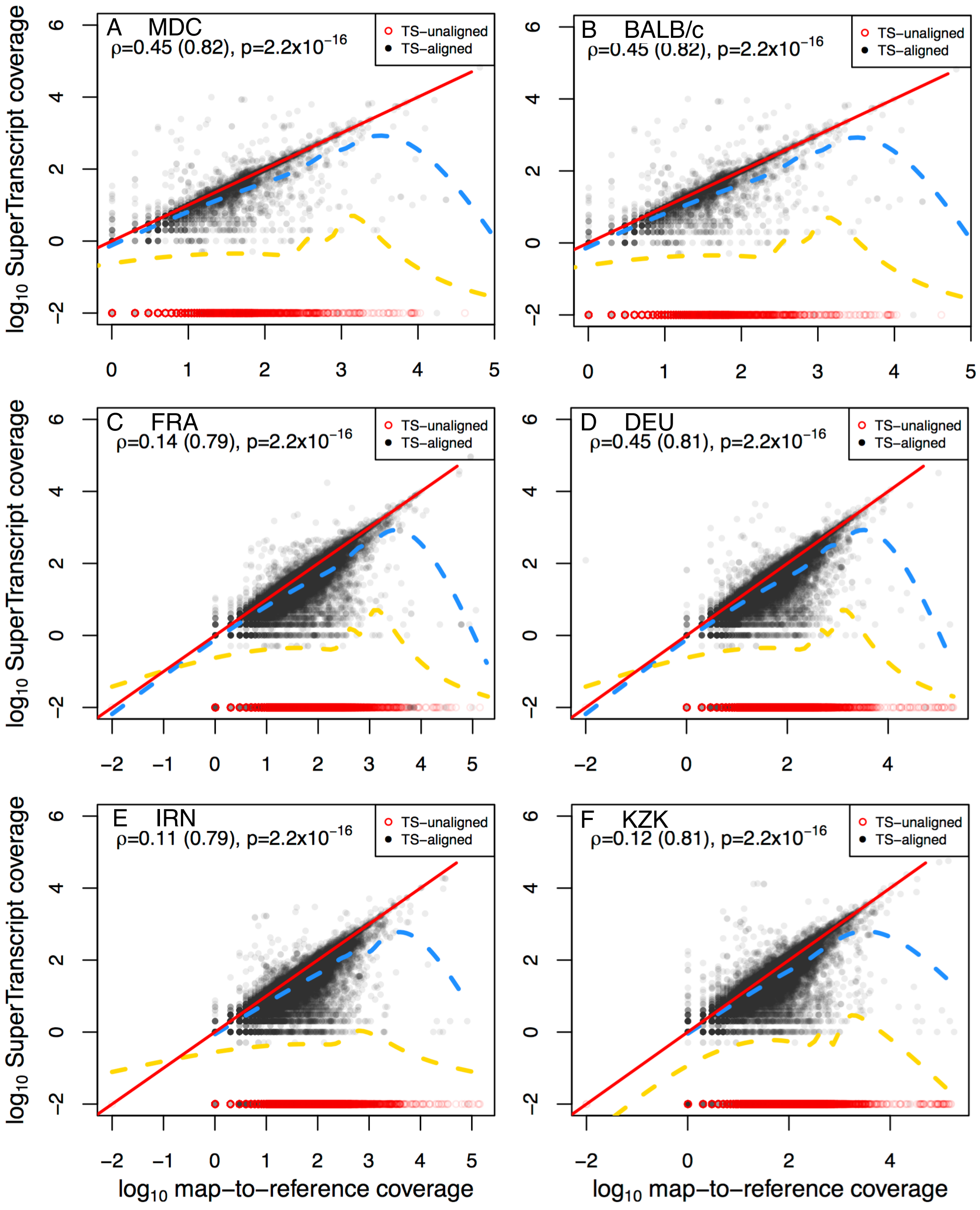


**Fig. S12.** Biplots and spearman's rank correlations of map-to-reference read depth vs. SuperTranscript read depth at positions where there are SuperTranscript (ST) false negatives (FN). Red circles are for genomic positions where there is no ST alignment such that read depth is, by definition, zero. Blue lines indicated the fitted, smoothed trend for only sites where there is a ST alignment, while yellow lines are for all FN sites. Prior to log-transformation, 0.01 is added to all values so that zero values are not omitted from the plots. Thus, log-transformed values of -2 indicate read depth of zero.

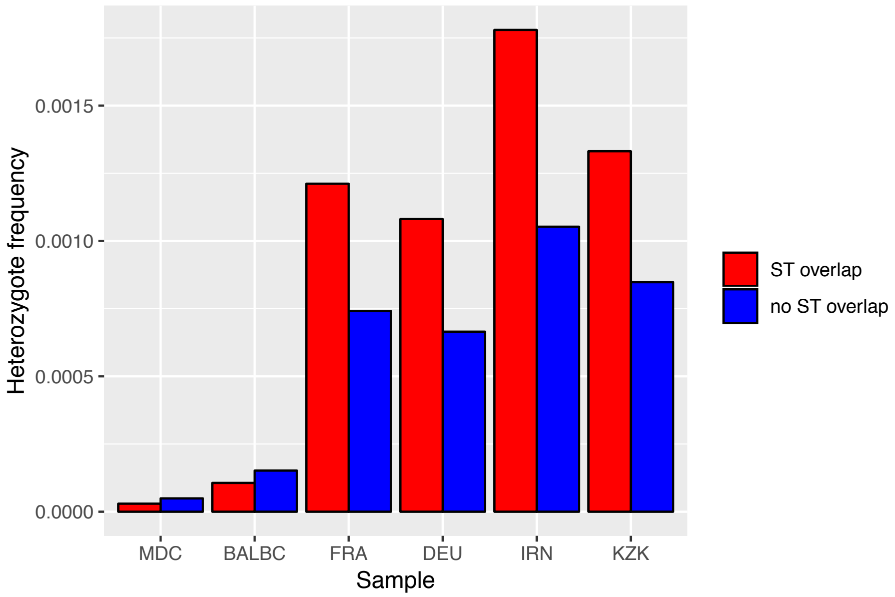


**Fig. S13.** Comparison of heterozygosity per base for biallelic SNV exonic variants derived from map-to-reference genotyping between regions with and and without overlapping SuperTranscripts.

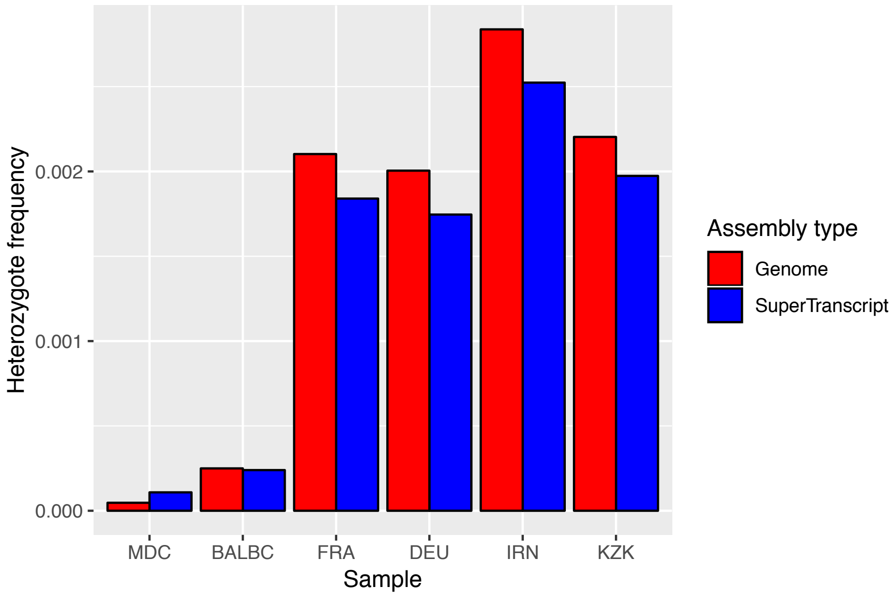


**Fig. S14.** Heterozygosity per base for all biallelic SNV variants called relative to the *Mus* reference genome using the GATK best practices pipeline versus those obtained from SuperTranscripts constructed from Trinity assemblies using scripts provided with the Trinity v. 2.6.5 package.

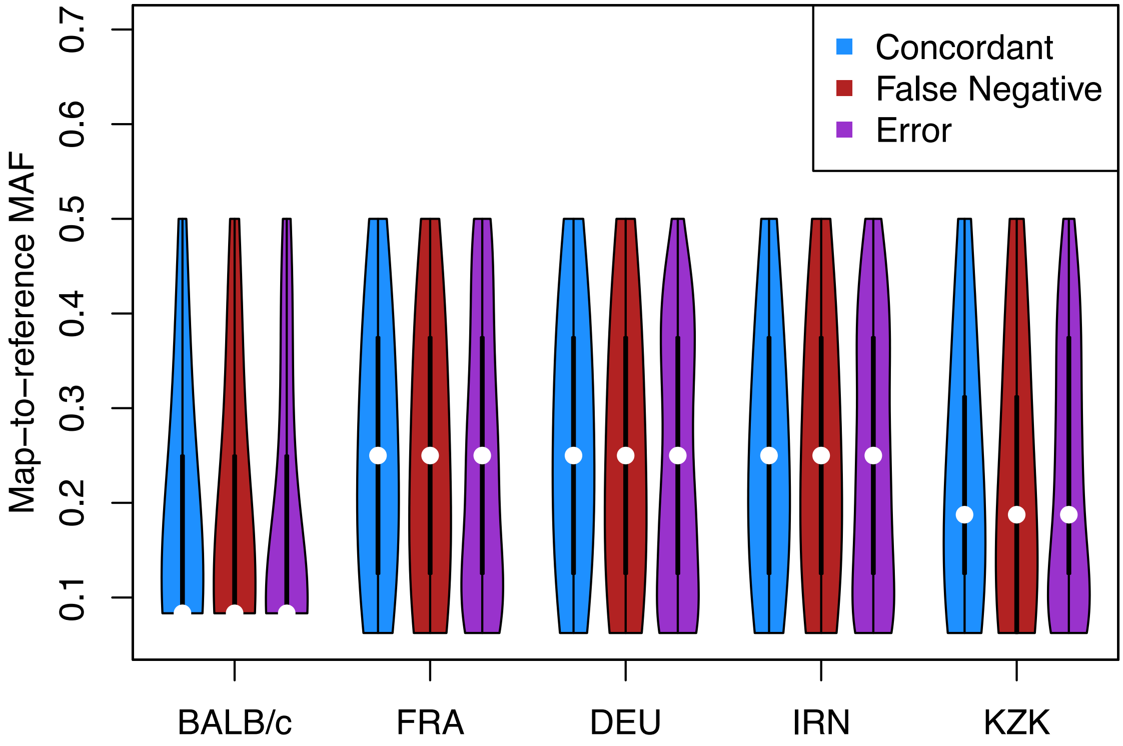


**Fig. S15.** Violin plots of map-to-reference (MR) minor allele frequencies at called heterozygous positions, as a function of SuperTranscript (ST) genotype calls at those positions that were either concordant (i.e. correct heterozygous calls), false negatives, or genotyping errors relative to MR.

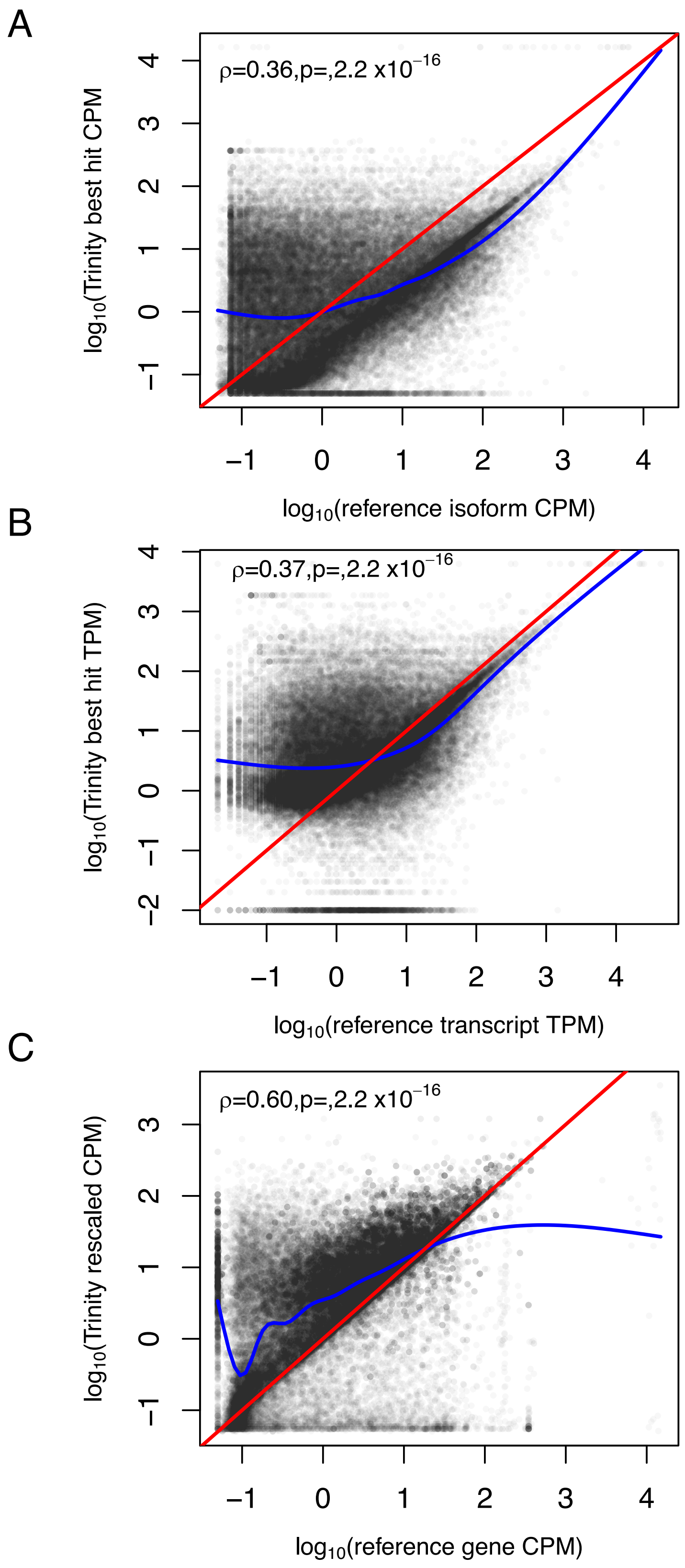


**Fig. S16.** For the FRA data set, comparison of expression between BLAT best hit Trinity contigs and reference annotations, in terms of (A) contig vs. reference isoform CPM, (B) contig vs. reference isoform CPM, and (C) contig rescaled CPM vs. reference gene CPM. The red line represents unity, the blue line is the smoothed fit to all of the data.

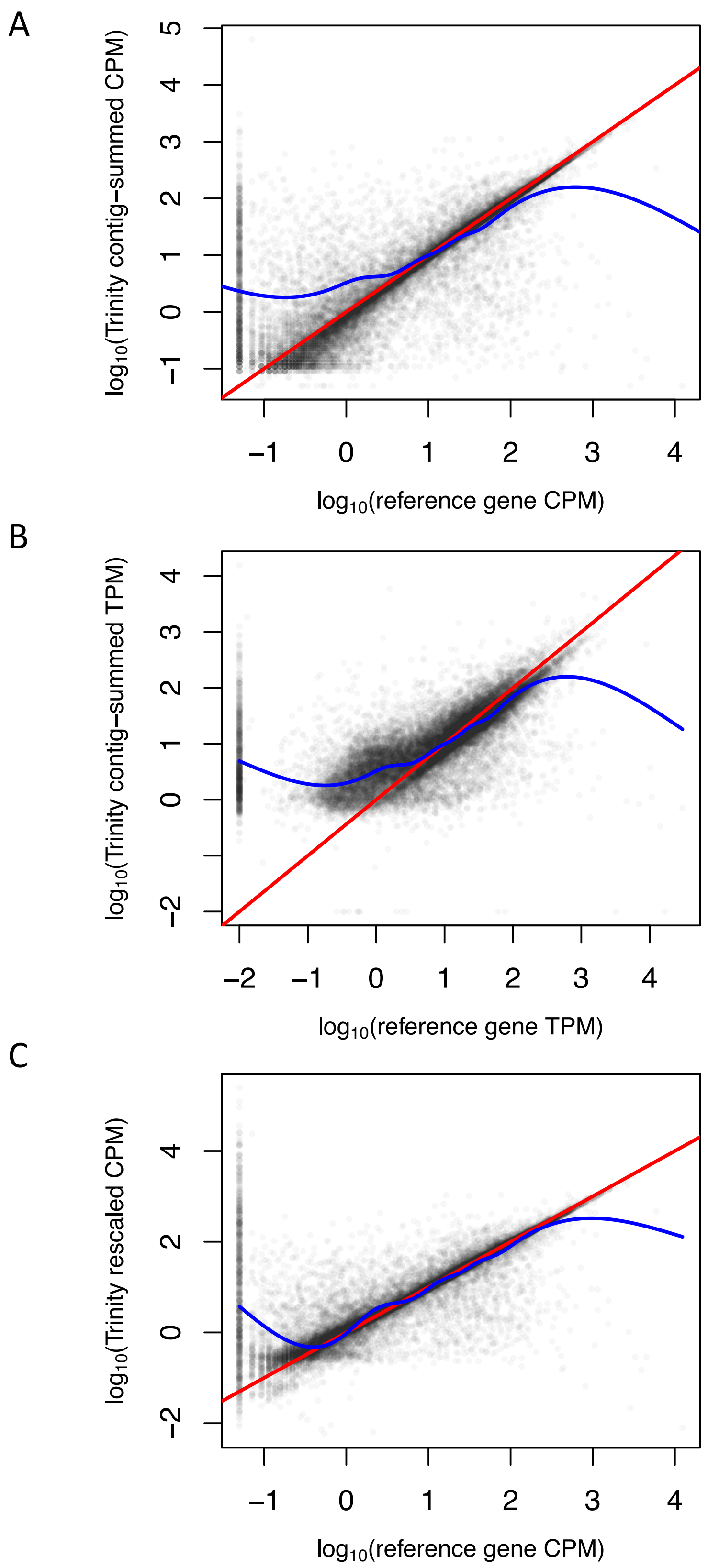


**Fig. S17.** Kallisto-derived comparisons of gene-level expression estimates between map-to-reference and summing abundance across Trinity contigs for the FRA sample for (A) CPM, (B) TPM, and (C) a corrected CPM for Trinity to account for differences in effective length. Spearman's rank correlations are, respectively, 0.86, 0.79, and 0.84.

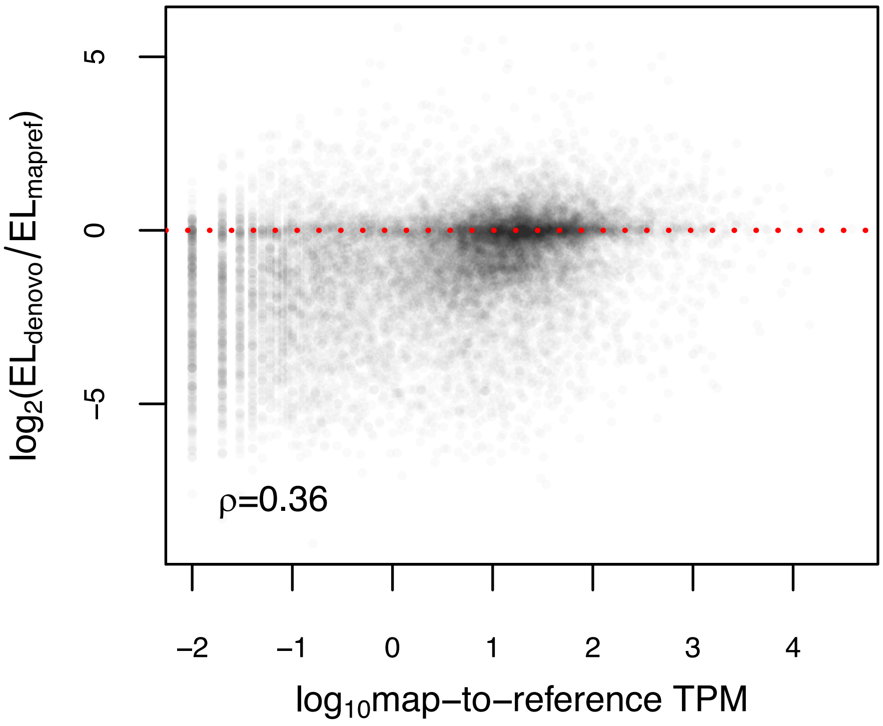


**Fig. S18.** For the FRA sample, log-fold change in effective length estimates for Trinity assembly vs. map-to-reference gene-level expression estimates as a function of reference expression in TPM units.

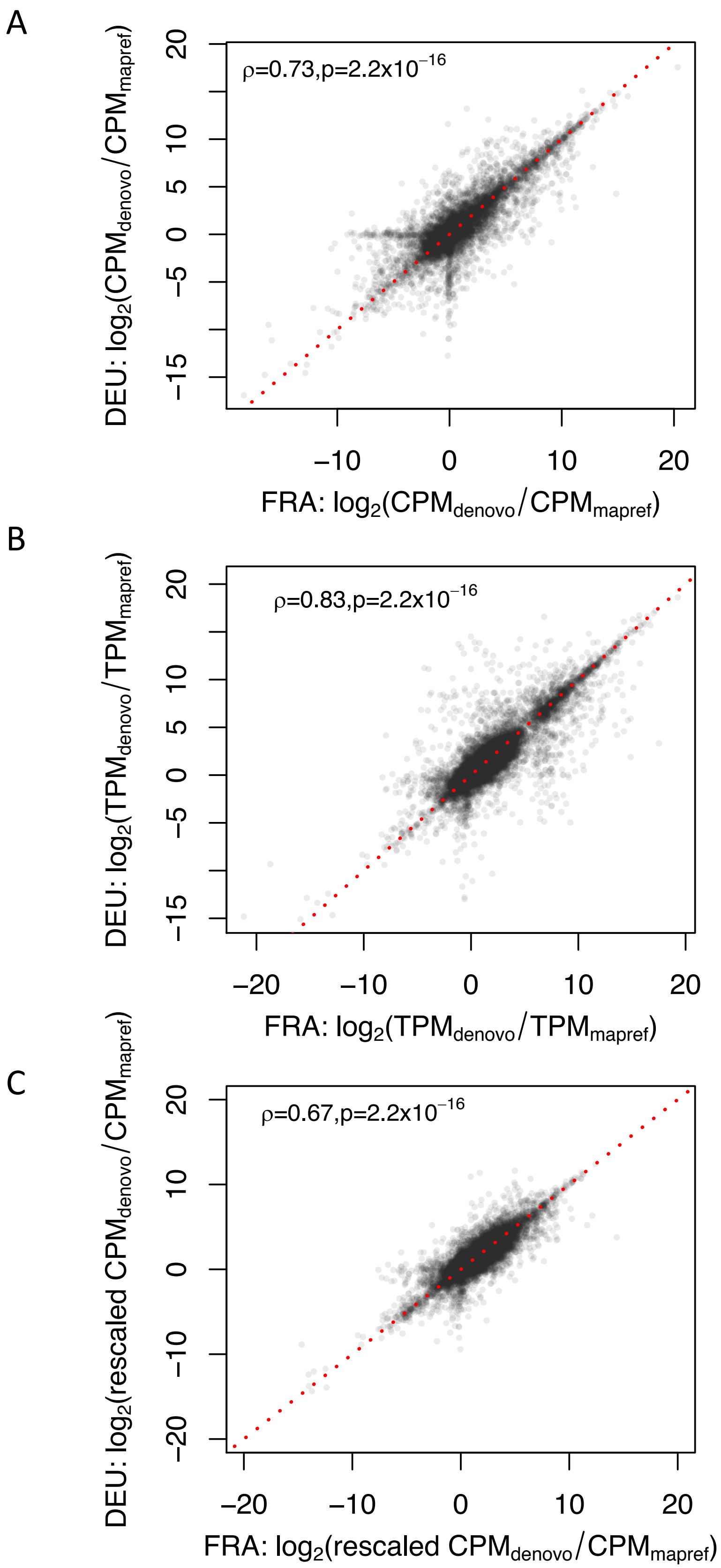


**Fig. S19.** Correlations in gene-level expression bias between the FRA and DEU samples, with bias measured as log-fold change between Trinity and map-to-reference based (A) CPM (B) TPM, and (C) rescaled CPM which corrects for differences in effective length estimates between the methods.
